# AMD1-driven MTA production sustains chemoresistance by rewiring PMN-MDSC immunosuppression

**DOI:** 10.64898/2026.11.02.763561

**Authors:** Yinan Wang, Wen Qiao, Bin Yu, Cegui Hu, Hao Li, Hanjun Li, Rongyu Zhang, Luying Li, Peng Li, Nannan Sha, Shijia Bao, Shanni Guo, Guiying Wei, Dongxi Xiang, Jun Gui

**Author notes:** These authors contributed equally (Yinan Wang, Wen Qiao, Bin Yu). Correspondence to: **Dr. Jun Gui**, State Key Laboratory of Systems Medicine for Cancer, Shanghai Cancer Institute, Ren Ji Hospital, Shanghai Jiao Tong University School of Medicine, Shanghai, 200127, China,; **Dr. Dongxi Xiang**, State Key Laboratory of Systems Medicine for Cancer, Shanghai Cancer Institute, Department of Biliary-Pancreatic Surgery, Department of Gastrointestinal Surgery, Renji Hospital, Shanghai Jiao Tong University School of Medicine, Shanghai, 200127, China.

## Abstract

Chemoresistance reflects both tumor cell-intrinsic adaptation and remodeling of the immune microenvironment. However, the mechanisms by which chemoresistant tumor cells orchestrate and sustain an immunosuppressive microenvironment remain poorly understood. Here, we demonstrate that chemoresistant tumors exhibit marked accumulation of polymorphonuclear myeloid-derived suppressor cells (PMN-MDSCs), which exert potent immunosuppressive effects on CD8^+^T cells and are critical for drug resistance. Metabolomic profiling reveals that resistant tumor cells release the metabolite 5’-methylthioadenosine (MTA). MTA enhances the immunosuppressive function of PMN-MDSCs by bolstering JAK-STAT3 signaling. Mechanistically, MTA binds to the protein tyrosine phosphatase PTPN2 to block its phosphatase activity, thereby potentiating activation of JAK-STAT3 signaling pathway. Moreover, we uncover that adenosylmethionine decarboxylase 1 (AMD1) is the key metabolic determinant of MTA accumulation in chemoresistant tumor cells. Genetic or pharmacological AMD1 inhibition significantly decreases MTA level, diminishes PMN-MDSC immunosuppressive function, and re-sensitizes the resistant tumors to chemotherapy and immunotherapy. Our findings identify MTA as an important metabolic link between chemoresistant tumor cells and PMN-MDSCs to sustain immunosuppression and therapeutic resistance. Targeting AMD1-dependent MTA production represents a valuable strategy for overcoming chemo-immunotherapy resistance.

## Introduction

Chemotherapy, either alone or in combination with immunotherapy, remains a cornerstone of treatment for many solid tumors, yet the emergence of acquired resistance often limits durable therapeutic responses. Although chemoresistance has traditionally been attributed to tumor-cell-intrinsic mechanisms, including enhanced drug efflux, increased DNA damage repair, evasion of apoptosis and acquisition of stem-like properties, these mechanisms are insufficient to fully account for treatment failure^1–3^. Resistant tumor cells also remodel the surrounding immune microenvironment, which in turn reinforces tumor cell survival and limits the efficacy of chemotherapy and immunotherapy^4,5^. Among the immune populations involved, polymorphonuclear myeloid-derived suppressor cells (PMN-MDSCs) are pathologically activated neutrophil-like myeloid cells that restrict antitumor immunity^6^. PMN-MDSCs suppress CD8^+^T cell proliferation and effector function through multiple mechanisms involving arginase 1 (ARG1), nitric oxide (NO), reactive oxygen species (ROS) and other immunoregulatory mediators^7,8^. Beyond their role in immune suppression, MDSCs can promote tumor-cell stemness and tumor propagation, thereby linking immune dysfunction to treatment-refractory tumor phenotypes^9^. Although PMN-MDSC accumulation has been associated with impaired responses to chemotherapy and immune checkpoint blockade, the tumor-derived signals that drive and sustain their suppressive state during acquired chemoresistance remain poorly defined.

Metabolites are increasingly recognized as active regulators of the tumor immune microenvironment ^10,11^. Under sustained selective pressure imposed by chemotherapy, tumor cells undergo metabolic adaptions that alter the production and release of metabolites, which in turn reshape the functional states of surrounding immune cells^12^. These metabolic alterations also contribute to chemotherapy resistance by generating a microenvironment that simultaneously enhances tumor cell survival and compromises immune-mediated tumor elimination^13,14^. Thus, metabolites constitute a critical molecular link connecting tumor metabolic reprogramming, immune suppression and chemoresistance. PMN-MDSCs are particularly sensitive to the metabolic conditions of the tumor microenvironment and can undergo profound functional change in response to hypoxia, nutrient availability and tumor-derived metabolites^15,16^. Defining how specific metabolites regulate PMN-MDSC behavior and their interactions with tumor cells may therefore provide new insights into the metabolic basis of therapeutic resistance.

In this study, we demonstrate that chemotherapy induces tumor metabolic adaptation that drives 5’-methylthioadenosine (MTA) accumulation, which directly reprograms PMN-MDSCs through fostering JAK-STAT3 signaling to establish an immunosuppressive niche and promote chemoresistance. MTA is a sulfur-containing nucleoside that arises primarily as a metabolic by-product of polyamine biosynthesis^17,18^. During the synthesis of spermidine and spermine, S-adenosylmethionine decarboxylase (AMD1) generates decarboxylated S-adenosylmethionine (dcSAM), which serves as an aminopropyl donor for polyamine formation and is concomitantly converted to MTA^19,20^. Under physiological conditions, MTA is rapidly metabolized by methylthioadenosine phosphorylase (MTAP) and incorporated into the methionine salvage pathway^21,22^. However, disruption of MTA metabolism, particularly through MTAP loss (∼10%), results in MTA accumulation and has been associated with profound alterations in antitumor immunity^23–25^.

In most MTAP-intact settings, here we show that sustained chemotherapy increases AMD1 expression in tumor cells, resulting in elevated MTA production. MTA directly binds to PTPN2 and perturbs its phosphatase activity on JAK–STAT3 signaling, thereby reinforcing the immunosuppressive activity of PMN-MDSCs, consequently impairing CD8^+^T cell anti-tumor immunity and supporting tumor-cell stemness and drug resistance. Genetic or pharmacological inhibition of AMD1 reduces MTA availability, eliminates PMN-MDSC-mediated immunosuppression and restores responsiveness to chemotherapy and anti-PD-1 treatment in resistant tumors. Our findings reveal a new metabolic–immune circuit through which chemoresistant tumor cells actively establish an immunosuppressive microenvironment that sustains drug resistance, positioning the AMD1-MTA–PMN-MDSCs axis as a potential therapeutic vulnerability for overcoming chemotherapy resistance.

## Results

### Chemoresistant tumors accumulate PMN-MDSCs and are refractory to chemo-immunotherapy

By continuous low-dose drug selection in *vitro*, we generated paclitaxel (PTX)-resistant breast cancer (4T1R) and cisplatin (CDDP)-resistant colorectal carcinoma (MC38R) tumor cell lines. Both resistant derivatives displayed higher half-maximal inhibitory concentrations (IC50 values) for their respective drugs compared with the corresponding parental cells (Extended Data Fig.1a). Following implantation into syngeneic mice, PTX or CDDP treatment significantly inhibited the growth of parental tumors but had little effect on their matched resistant tumors, confirming that the resistant phenotypes were maintained in *vivo* (Fig. 1a and Extended Data Fig.1b). Of note, the resistant tumors were also refractory to anti-PD-1 therapy, either alone or in combination with chemotherapy (Fig.1b and Extended Data Fig.1c), indicating that the resistance tumors might remodel the tumor immune microenvironment. We analyzed the immune profiling of tumors from both models and found that PMN-MDSCs were remarkably accumulated in the resistant tumors (Fig.1c). We next examined whether the same immune features were evident in human tumors. By deconvoluting transcriptomic data of three independent cohorts of breast cancer patients received with neo-adjuvant chemotherapy, we observed higher PMN-MDSC gene scores, but lower CD8^+^T cell scores in the non-responders when compared to the responders (Fig. S1d). Moreover, high PMN-MDSC signatures were associated with shorter survival rate (Extended Data Fig. 1e). We further analyzed the public single-cell RNA-sequencing data from colorectal cancer patients (Pembrolizumab + Capecitabine + Oxaliplatin) and breast cancer patients (Pembrolizumab + Carboplatin + Paclitaxel + Anthracycline) with chemo-immunotherapy treatment. Similarly, PMN-MDSC fraction was greatly increased in the resistant patients (Fig. 1d and Extended Data Fig.1f). Cell–cell communication analysis showed stronger predicted interactions of PMN-MDSCs with tumor cells and T cells in resistant patients (Fig. 1e). Meanwhile, PMN-MDSCs from resistant patients expressed higher levels of the immunosuppressive genes, whereas their T cells showed reduced expression of activation and cytotoxicity-associated genes (Fig. 1f-g and Extended Data Fig.1g-h). Indeed, tumor cells from resistant patients displayed higher expression of stemness, adhesion and resistance-associated genes (Fig. 1h and Extended Data Fig.1i). Collectively, all these data suggest that chemoresistant tumors exhibit an immunosuppressive microenvironment, evidenced by a PMN-MDSC-rich and T cell-poor immune state.

**Fig. 1.**
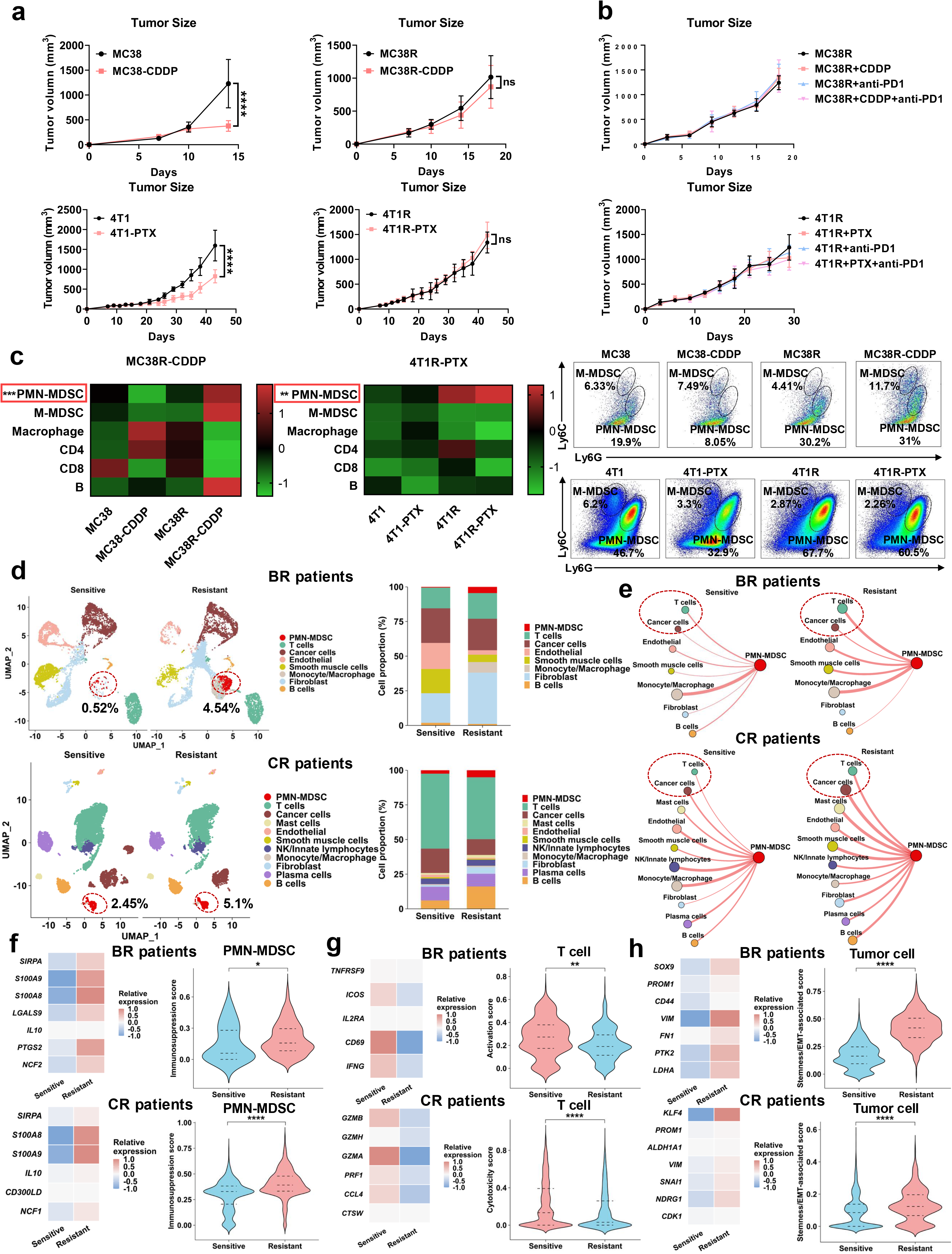
Chemoresistant tumors accumulate PMN-MDSCs and exhibit an immunosuppressive microenvironment. **(a)**Tumor growth of parental MC38 and 4T1 tumors and their chemoresistant counterparts MC38R and 4T1R, with or without treatment with the corresponding chemotherapeutic agents cisplatin (CDDP) or paclitaxel (PTX). n = 4-5 mice/group. Data are shown as mean ± SEM; repeated-measures two-way ANOVA followed by Sidak’s multiple-comparisons test. **(b)** Tumor growth of MC38R and 4T1R tumors treated with vehicle, chemotherapy, anti-PD-1, or combined chemotherapy and anti-PD-1. n = 4–5 mice/group. Data are shown as mean ± SEM; repeated-measures two-way ANOVA followed by Sidak’s multiple-comparisons test. **(c)** Heatmaps showing the relative abundance of tumor-infiltrating immune-cell populations in parental and chemoresistant MC38 and 4T1 tumors with or without chemotherapy. Representative flow cytometry plots show the frequencies of M-MDSCs and PMN-MDSCs in the indicated tumor groups. n=4-5 mice/group. **(d)** UMAP plots and relative proportions of different cell populations in treatment-sensitive and treatment-resistant breast cancer (GSE302453; top) and colorectal cancer (GSE236581; bottom) patient samples analyzed by scRNA-seq. **(e)** Predicted cell–cell communication networks among cell populations in treatment-sensitive and treatment-resistant breast cancer (top) and colorectal cancer (bottom) samples. PMN-MDSC-centered interactions are highlighted. **(f)** Expression heatmaps of PMN-MDSC immunosuppression-associated genes and corresponding UCell signature scores in treatment-sensitive and treatment-resistant breast cancer (top) and colorectal cancer (bottom) samples. **(g)** Expression heatmaps of T cell activation/cytotoxicity-associated genes and corresponding UCell signature scores. **(h)** Expression heatmaps of tumor stemness-, adhesion-, and resistance-associated genes and corresponding UCell signature scores in malignant cells.

### PMN-MDSCs from resistant tumors suppress CD8^+^T cell function and sustain chemoresistance

To determine whether the resistant tumor cells directly promoted the suppressive effects of PMN-MDSCs, we isolated Ly6G^+^ neutrophils from mouse bone marrow and cultured them for 5 days in the presence of granulocyte–macrophage colony-stimulating factor (GM-CSF) and tumor cell-conditioned medium (TCM). The resulting PMN-MDSCs were then co-cultured with OT-1 CD8^+^T cells for 72 h (Fig. 2a). In both the 4T1 and MC38 models, PMN-MDSCs generated with resistant-cell TCM more potently inhibited T-cell proliferation and cytotoxic function when compared to those generated with parental-cell TCM (Fig. 2b-c), suggesting that soluble factors released by resistant tumor cells induced a more suppressive PMN-MDSC phenotype.

**Fig. 2.**
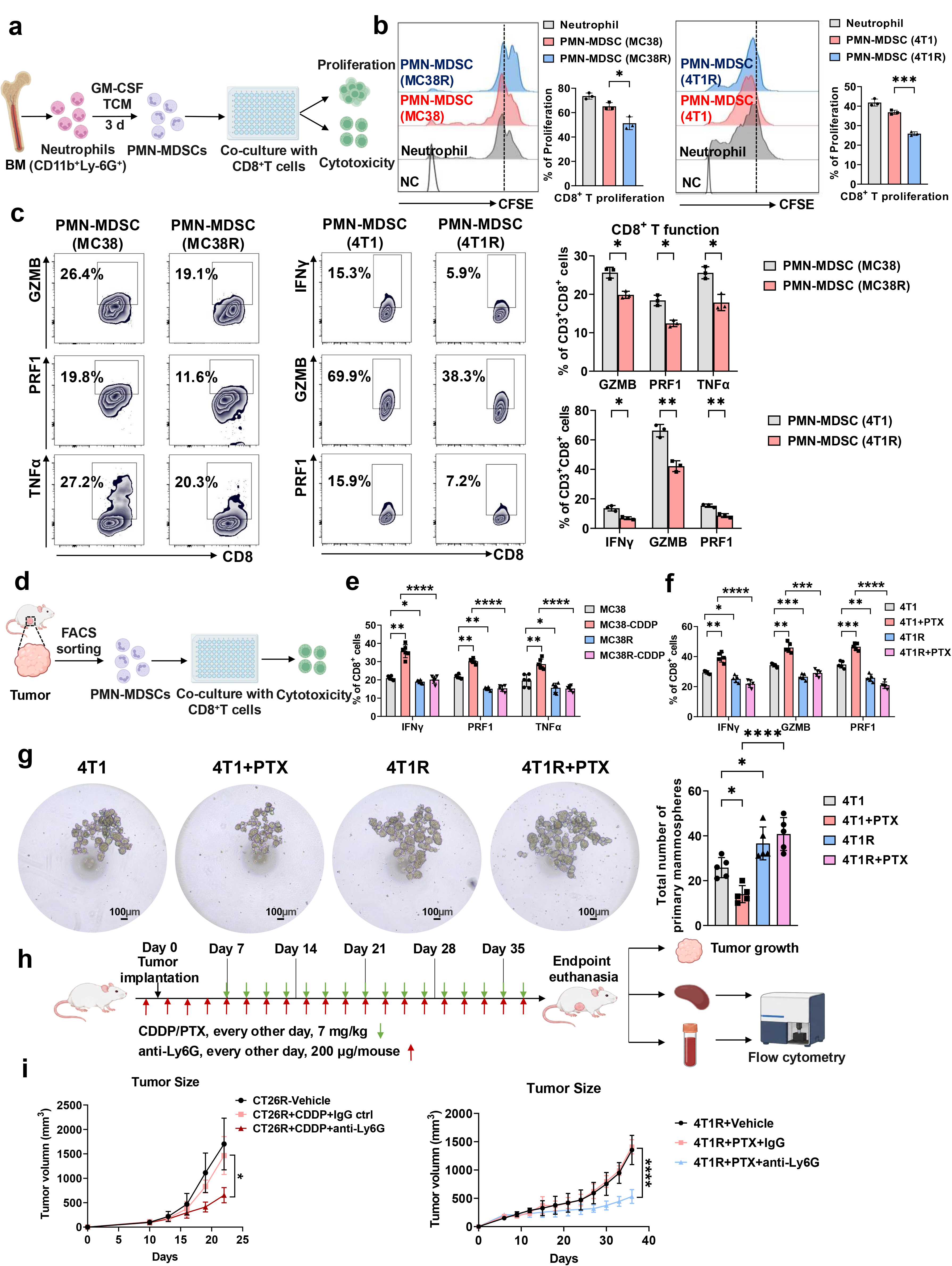
PMN-MDSCs from chemoresistant tumors suppress CD8^+^T cell function and sustain chemoresistance. **(a)** Schematic illustration for the generation and functional assessment of TCM-induced PMN-MDSCs. **(b)** Representative CFSE profiles and quantification of CD8^+^T cell proliferation following co-culture with neutrophils or PMN-MDSCs induced by parental or chemoresistant MC38 and 4T1 TCM. n = 3 biological replicates. Data are shown as mean ± SEM; one-way ANOVA followed by Sidak’s multiple-comparisons test. **(c)** Representative flow cytometry plots and quantification of GZMB, PRF1, TNF-α, and IFN-γ expression in CD8^+^T cells following co-culture with PMN-MDSCs induced by parental or chemoresistant MC38 and 4T1 TCM. n = 3 biological replicates/group. Data are shown as mean ± SEM; two-way ANOVA followed by Sidak’s multiple-comparisons test. **(d)** Schematic illustration for the ex vivo functional assay using PMN-MDSCs isolated from tumors, followed by co-culture with CD8^+^T cells. **(e-f)** Cytotoxic effector molecule expression in CD8^+^T cells following co-culture with PMN-MDSCs isolated from parental or chemoresistant MC38 **(e)** and 4T1 **(f)** tumors with or without chemotherapy. n = 5 biological replicates/group. Data are shown as mean ± SEM; two-way ANOVA followed by Sidak’s multiple-comparisons test. **(g)** Representative images and quantification of primary mammosphere formation following co-culture with PMN-MDSCs isolated from the indicated tumor groups. n = 5 biological replicates/group; one-way ANOVA followed by Sidak’s multiple-comparisons test. Scale bars, 100 μm. **(h)** Schematic illustration for Ly6G^+^ cell depletion during chemotherapy treatment of chemoresistant tumors. **(i)** Tumor growth of CT26R and 4T1R tumors treated with vehicle or chemotherapy together with control IgG or anti-Ly6G antibody. n = 5-6 mice/group. Data are shown as mean ± SEM; repeated-measures two-way ANOVA followed by Sidak’s multiple-comparisons test.

We next isolated PMN-MDSCs directly from parental or resistant tumors with or without chemotherapy and tested their suppressive activity ex vivo (Fig. 2d). Notably, PMN-MDSCs from chemotherapy-treated parental tumors were less suppressive than those from untreated parental tumors, as indicated by greater T cell proliferation and cytotoxic effector expression after co-culture. By contrast, PMN-MDSCs from resistant tumors remained strongly suppressive effects on T cells regardless of chemotherapy exposure (Fig. 2e-f and Extended Data Fig. 2a-b). In parallel, PMN-MDSCs isolated from resistant tumors increased 4T1 tumor-cell sphere formation relative to those from parental tumors (Fig. 2g), indicating that resistant-tumor-associated PMN-MDSCs induced both T cell dysfunction and tumor cell propagation.

To validate the critical role of PMN-MDSCs in chemoresistance, we depleted PMN-MDSCs by injecting anti-Ly-6G antibody in the mice bearing resistant tumors (Fig. 2h). Anti-Ly6G treatment substantially depleted circulating and splenic Ly6G^+^ cells (Extended Data Fig. 2c). Importantly, this depletion apparently restored chemotherapy-mediated tumor control and reduced resistant-tumor burdens (Fig. 2i and Extended Data Fig. 2d). These data suggest that Ly6G^+^ neutrophils/PMN-MDSCs are essential for mediating chemoresistance.

### MTA accumulates in resistant tumors and enhances PMN-MDSC immunosuppressive activity

To identify what tumor-derived factor is responsible for the enhanced PMN-MDSC immunosuppression, we separated parental and resistant TCM into molecular-mass fractions using sequential ultrafiltration. Among the fractions tested in both 4T1 and MC38 TCM, we noticed that <3kDa fraction from resistant-cell TCM induced the strongest suppressive landscape than the other fractions, as reflected by robustly increasing expression of immunosuppressive markers in PMN-MDSCs, including *iNOS*, *Arg1*, *S100a8* and *S100a9* (Extended Data Fig. 3a-b), and prominent suppression on T cell proliferation (Extended Data Fig. 3c). As this fraction is mainly small-molecule metabolites, we performed untargeted metabolomic profiling of parental and resistant TCM. Intriguingly, we found that MTA was significantly increased both in 4T1R and MC38R-conditioned media (Fig. 3a). Targeted measurements further confirmed higher MTA levels in resistant-cell TCM and resistant-tumor interstitial fluid (TIF) in contrast to those from sensitive tumors (Fig. 3b).

**Fig. 3.**
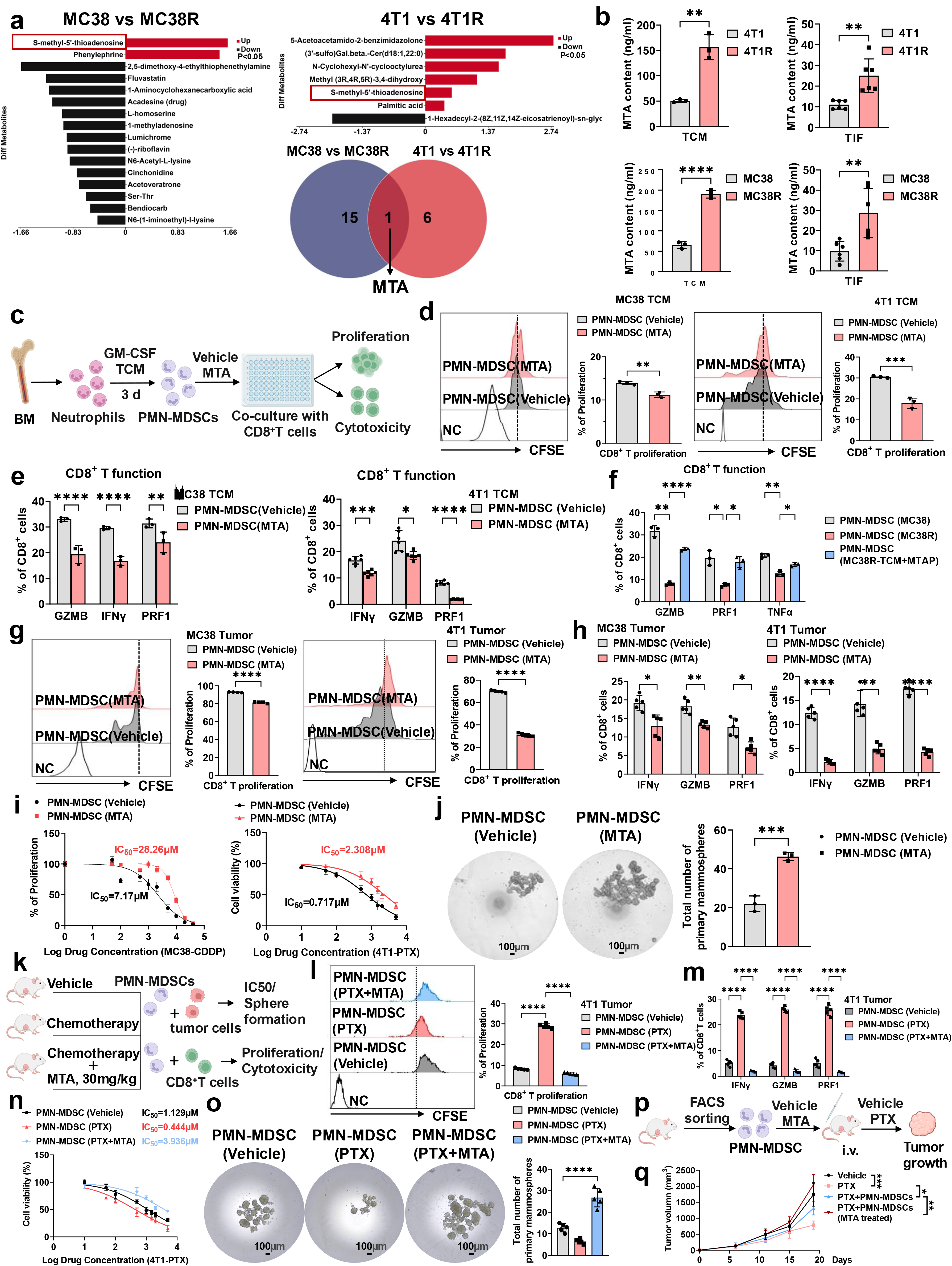
MTA accumulates in chemoresistant tumors and enhances PMN-MDSC immunosuppressive activity. **(a)** Untargeted metabolomic profiling of TCM from parental and chemoresistant MC38 and 4T1 cells. Differential metabolites are shown, and the Venn diagram identifies MTA as a metabolite commonly increased in both chemoresistant models. n = 3 independent samples/group. **(b)** Targeted quantification of MTA in TCM (n = 3 biological replicates/group) and TIF (n = 5 biological replicates/group) derived from parental and chemoresistant 4T1 and MC38 tumors. Data are shown as mean ± SEM; two-tailed unpaired Student’s t test. **(c)** Schematic illustration for the functional assessment of TCM-induced PMN-MDSCs with MTA treatment. **(d)** Representative CFSE profiles and quantification of CD8^+^T cell proliferation following co-culture with vehicle-or MTA-treated PMN-MDSCs. n = 3 biological replicates/group; two-tailed unpaired Student’s t test. **(e)** Quantification of GZMB, IFN-γ, and PRF1 expression in CD8^+^T cells following co-culture with vehicle-or MTA-treated PMN-MDSCs induced by MC38 or 4T1 TCM. n = 3 biological replicates/group. Data are shown as mean ± SEM; two-way ANOVA followed by Sidak’s multiple-comparisons test. **(f)** Cytotoxic effector molecule expression in CD8^+^T cells following co-culture with PMN-MDSCs induced by MC38, MC38R, or MC38R TCM added with MTAP. n = 3 biological replicates/group; two-way ANOVA followed by Sidak’s multiple-comparisons test. **(g)** Representative CFSE profiles and quantification of CD8^+^T cell proliferation following co-culture with vehicle-or MTA-treated PMN-MDSCs isolated from tumors. n = 5 biological replicates/group; two-tailed unpaired Student’s t test. **(h)** Quantification of IFN-γ, GZMB, and PRF1 expression in CD8^+^T cells following co-culture with vehicle-or MTA-treated PMN-MDSCs isolated from tumors. n = 5 biological replicates/group; two-way ANOVA followed by Sidak’s multiple-comparisons test. **(i)** Chemotherapeutic drug-response curves of MC38 and 4T1 tumor cells following co-culture with vehicle-or MTA-treated PMN-MDSCs. n = 3 biological replicates/group. Data are shown as mean ± SEM; two-way ANOVA. **(j)** Representative images and quantification of primary mammosphere formation following co-culture with vehicle-or MTA-treated PMN-MDSCs. n = 3 biological replicates/group; two-tailed unpaired Student’s t test. Scale bars, 100 μm. **(k)** Schematic illustration for the ex vivo functional assessment of PMN-MDSCs isolated from 4T1 tumor-bearing mice treated with vehicle, PTX, or PTX plus MTA. **(l)** Representative CFSE profiles and quantification of CD8^+^T cell proliferation following co-culture with PMN-MDSCs isolated from the indicated treatment groups. n = 5 mice/group; one-way ANOVA followed by Sidak’s multiple-comparisons test. **(m)** Quantification of IFN-γ, GZMB, and PRF1 expression in CD8^+^T cells following co-culture with PMN-MDSCs isolated from the indicated treatment groups. n = 5 mice/group; two-way ANOVA followed by Sidak’s multiple-comparisons test **(n)** PTX-response curves of 4T1 tumor cells following co-culture with PMN-MDSCs isolated from the indicated treatment groups. n = 5 mice/group; two-way ANOVA. **(o)** Representative images and quantification of primary mammosphere formation following co-culture with PMN-MDSCs isolated from the indicated treatment groups. N = 5 mice/group; one-way ANOVA followed by Sidak’s multiple-comparisons test. Scale bars, 100 μm. **(p)** Schematic illustration for the adoptive transfer of vehicle-or MTA-treated PMN-MDSCs into 4T1 tumor-bearing mice receiving PTX treatment. **(q)** Tumor growth following adoptive transfer of the indicated PMN-MDSCs. n = 5 mice/group. Data are shown as mean ± SEM; repeated-measures two-way ANOVA followed by Sidak’s multiple-comparisons test.

We next assessed whether MTA was sufficient to reinforce PMN-MDSC-mediated suppression. Bone-marrow neutrophils were differentiated with parental-cell TCM, then exposed to vehicle or MTA for 24 h before co-culture with CD8^+^T cells (Fig. 3c). Obviously, MTA-treated PMN-MDSCs more strongly inhibited T cell proliferation and reduced expression of cytotoxic molecules including IFN-γ, Granzyme B and Perforin-1 (Fig. 3d-e and Extended Data Fig. 3d). Conversely, enzymatic depletion of extracellular MTA from resistant-cell TCM using MTAP attenuated the suppressive activity of the resulting PMN-MDSCs (Fig. 3f and Extended Data Fig. 3e). Likewise, MTA enhanced the suppressive activity of PMN-MDSCs isolated from MC38 and 4T1 tumors, resulting in stronger inhibitory effects on T cell proliferation and cytotoxic function (Fig. 3g-h and Extended Data Fig. 3f). Moreover, MTA-treated PMN-MDSCs increased the IC50 doses of chemotherapeutic agents in tumor cells (Fig.3i) and promoted tumor-sphere formation (Fig.3j and Extended Data Fig.3g), thereby intensifying chemoresistance. We then supplemented MTA in mice bearing 4T1 tumors (sensitive) during PTX treatment (Fig. 3k). We found that MTA completely abolished the anti-tumor effect of PTX treatment (Extended Data Fig.3h-i). PMN-MDSCs isolated from mice receiving combined PTX and MTA treatment exhibited substantially greater suppression of T cell proliferation and effector functions than PMN-MDSCs isolated from mice treated with PTX alone (Fig. 3l-m and Extended Data Fig. 3j). Furthermore, these PMN-MDSCs increased the PTX IC50 values of tumor cells and enhanced tumor-sphere formation (Fig. 3n-o), promoting tumor cell drug resistance. Finally, we adoptively transferred vehicle or MTA-treated PMN-MDSCs into tumor-bearing mice undergoing PTX treatment (Fig. 3p). Indeed, PMN-MDSCs transfer attenuated PTX-mediated tumor control. Apparently, MTA-pretreated PMN-MDSCs completely abrogated the therapeutic efficacy of PTX (Fig. 3q), leading to pronounced chemoresistance. Taken together, these results suggest that resistant tumor cell-derived MTA reinforced PMN-MDSC-mediated immunosuppression to sustain chemoresistance.

### MTA activates JAK–STAT3 signaling in PMN-MDSCs by blocking PTPN2 phosphatase activity

To explore the potential mechanism underlying MTA enhances the immunosuppressive activity of PMN-MDSCs, we performed RNA sequencing of vehicle- and MTA-treated PMN-MDSCs. Gene set enrichment analysis (GSEA) revealed that immune-active pathways, including antigen presentation and T cell activation, were significantly downregulated in MTA-treated PMN-MDSCs (Extended Data Fig. 4a). Of note, JAK–STAT3 signaling was among the top 20 enriched pathways upregulated upon MTA exposure (Fig. 4a). JAK-STAT3 signaling has previously been implicated in PMN-MDSC mediated immunosuppression^26,27^, as its activation promotes the transcription of the immunosuppressive genes, such as *Arg1*, *S100a8* and *S100a9* (Fig. 4b). Accordingly, immunoblotting demonstrated increased phosphorylation of JAK1 and STAT3 in PMN-MDSCs following MTA treatment (Fig. 4c). Moreover, pharmacological inhibition of STAT3 significantly attenuated the expression of immunosuppressive genes induced by MTA (Fig. 4d), suggesting that JAK–STAT3 signaling contributes to MTA-mediated reprogramming of PMN-MDSCs.

**Fig. 4.**
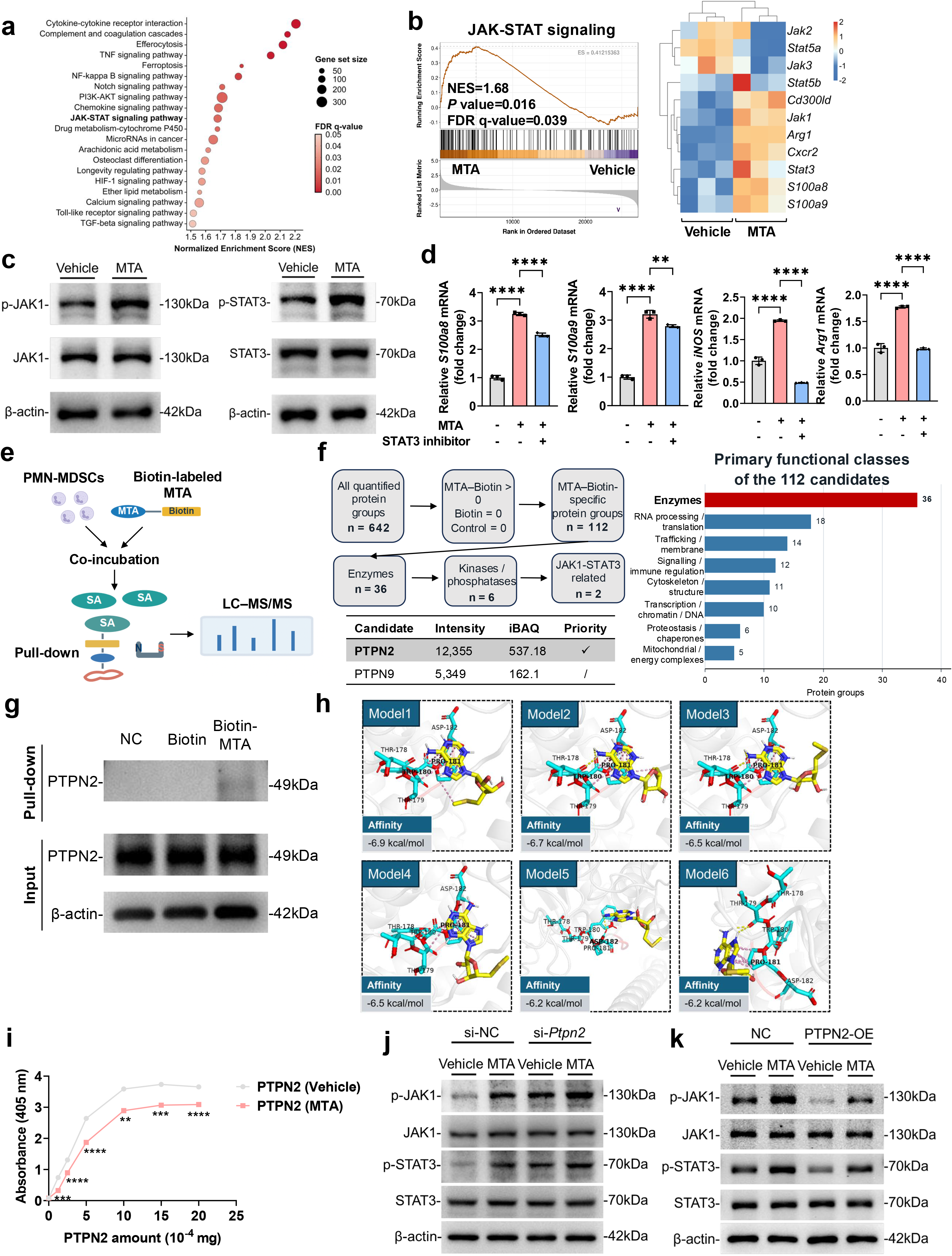
MTA activates JAK–STAT3 signaling in PMN-MDSCs by inhibiting PTPN2 phosphatase activity. **(a)** Ranking of signaling pathways enriched in MTA-treated PMN-MDSCs based on RNA-seq analysis. n = 3 biological replicates/group. **(b)** GSEA analysis of JAK–STAT signaling in MTA-treated PMN-MDSCs and heatmap of representative pathway-associated genes. n = 3 biological replicates/group. **(c)** Immunoblot analysis of phosphorylated and total JAK1 and STAT3 in vehicle-or MTA-treated PMN-MDSCs. Blots are representative of three independent experiments. **(d)** Relative expression of *S100a8, S100a9, Nos2*, and *Arg1* in PMN-MDSCs treated with vehicle or MTA in the presence or absence of STAT3 inhibitor Stattic. n = 3 biological replicates/group. Data are shown as mean ± SEM; two-way ANOVA followed by Sidak’s multiple-comparisons test. **(e)** Schematic illustration for the biotin–MTA affinity purification and LC–MS/MS workflow. **(f)** Filtering and functional classification of candidate MTA-associated proteins identified by biotin–MTA affinity purification and LC–MS/MS. Intensity and iBAQ values of PTPN2 and PTPN9 are shown. Three biological samples from each group were pooled before LC–MS/MS analysis, resulting in one pooled sample per group for proteomic profiling. **(g)** Streptavidin pull-down followed by immunoblotting for PTPN2. Blots are representative of three independent pull-down experiments. **(h)** Six predicted molecular docking poses of MTA with PTPN2 and corresponding Vina scores. n = 6 computational docking poses. **(i)** Recombinant PTPN2 phosphatase activity at increasing protein concentrations in the presence or absence of MTA. n = 3 independent experiments. Data are shown as mean ± SEM; two-way ANOVA followed by Sidak’s multiple-comparisons test. **(j)** Immunoblot analysis of phosphorylated and total JAK1 and STAT3 in control or *Ptpn2*-silenced PMN-MDSCs treated with vehicle or MTA. Blots are representative of three independent experiments. **(k)** Immunoblot analysis of phosphorylated and total JAK1 and STAT3 in control or PTPN2-overexpressing PMN-MDSCs treated with vehicle or MTA. Blots are representative of three independent experiments.

Next, we sought to identify the direct engaging molecule responsible for MTA-induced activation of JAK–STAT3 signaling. To this end, we synthesized biotin-labeled MTA (Extended Data Fig. 4b). PMN-MDSCs were incubated with biotin-labeled MTA or control biotin, and the lysates were pull down by Streptavidin, subsequently subjected to mass spectrometry (Fig. 4e). In total, we identified 112 proteins that may specifically interact with MTA, most of which are enzymes. Among these enzyme candidates, we noticed two enzymes, protein tyrosine phosphatase non-receptor type 2 (PTPN2) and PTPN9, which are well recognized as the negative regulators of JAK– STAT signaling via direct dephosphorylation of JAKs and STATs^28,29^. Notably, PTPN2 exhibited a stronger binding affinity for MTA than PTPN9 (Fig. 4f and Extended Data Fig.4c). Indeed, immunoprecipitation followed by immunoblotting validated the interaction of MTA with PTPN2 (Fig. 4g). By molecular docking, we predicted the binding sites of PTPN2 with MTA. Multiple models all pointed that MTA-binding poses were on the PTPN2 WPD-loop region (TRP180, PRO181, ASP182) (Fig. 4h and Extended Data Fig. 4d-e), which is critical for efficient substrate dephosphorylation^30^. In vitro assay using recombinant PTPN2 further revealed that MTA treatment robustly decreased its phosphatase activity (Fig. 4i). Consistent with the role of PTPN2 as a restraint on JAK–STAT signaling, *Ptpn2* knockdown increased JAK1 and STAT3 phosphorylation in PMN-MDSCs, and MTA treatment slightly enhanced its activation (Fig. 4j and Extended Data Fig. 4f). Conversely, PTPN2 overexpression in PMN-MDSCs markedly suppressed JAK1 and STAT3 phosphorylation, although MTA treatment substantially increased its activation when compared to the vehicle treatment (Fig.4k and Extended Data Fig. 4g). Collectively, all these data suggest that MTA binding disrupts PTPN2-mediated negative control of JAK–STAT3 signaling, thereby promoting and sustaining the immunosuppressive state of PMN-MDSCs.

### AMD1-driven MTA production by chemoresistant tumor cells promotes the immunosuppressive activity of PMN-MDSCs

Next, we sought to determine the mechanism underlying MTA accumulation in chemoresistant tumor cells. MTA is generated as a by-product of polyamine biosynthesis, a metabolic pathway regulated by three key enzymes: AMD1, spermidine synthase (SRM) and spermine synthase (SMS) (Fig. 5a). MTA can be metabolized through MTAP in the downstream to maintain intracellular MTA homeostasis. Numerous studies have demonstrated that MTAP deletion in tumor leads to aberrant accumulation of MTA^21–24^. However, we found that MTAP was readily expressed in both MC38 and 4T1 tumor cells, and its expression exhibited no difference between parental and resistant tumor cells (Extended Data Fig. 5a), suggesting that MTA accumulation in chemoresistant tumor cells was unlikely to result from impaired MTAP expression. We therefore assumed that the upstream synthetic enzymes might account for the increased MTA production and the enhanced immunosuppressive activity of PMN-MDSCs. We individually silenced *Amd1*, *Srm* and *Sms* in the resistant tumor cells and used the resulting TCM to induce PMN-MDSCs. Among these perturbations, *Amd1* silencing most pronouncedly reduced the suppressive activity of TCM-conditioned PMN-MDSCs, as shown by improved CD8^+^T cell effector function after co-culture (Extended Data Fig. 5b). Indeed, AMD1 expression was significantly upregulated in the resistant tumor cells when compared to their parental counterparts (Fig. 5b). We therefore generated stable *Amd1*-knockdown resistant tumor cells and confirmed efficient reduction of AMD1 expression by RT–qPCR and immunoblotting (Extended Data Fig.5c-d). Obviously, MTA levels were markedly decreased in the TCM derived from AMD1-knockdown resistant tumor cells, indicating that AMD1 is required for MTA production in these cells (Fig. 5c). In addition, we noticed that the AMD1-involved metabolites spermidine and spermine had minimal effects on the expression of immunosuppressive markers in PMN-MDSCs (Extended Data Fig. 5e).

**Fig. 5.**
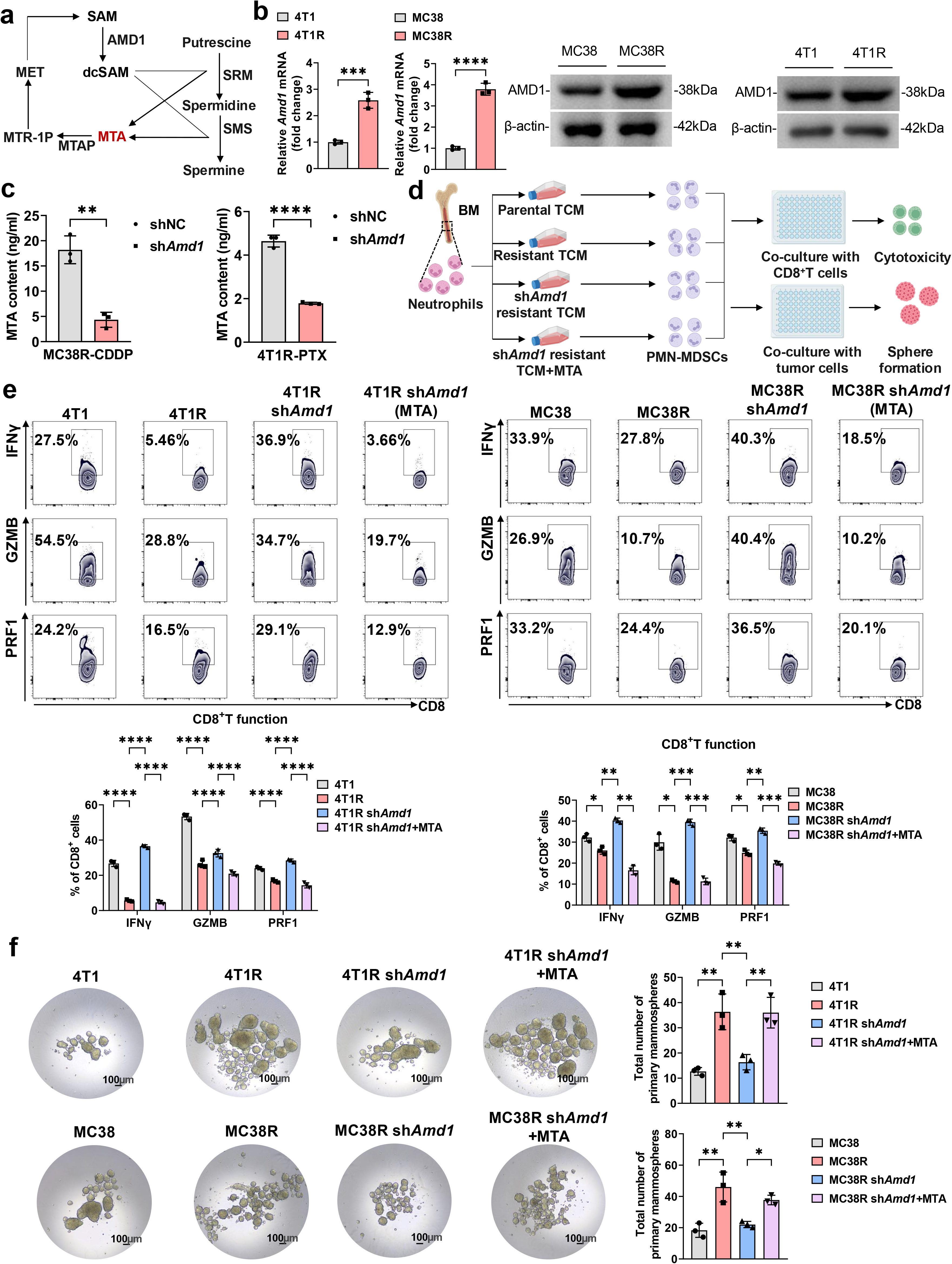
AMD1-driven MTA production by chemoresistant tumor cells promotes PMN-MDSC immunosuppressive activity. **(a)** Schematic illustration for polyamine biosynthesis and MTA metabolism. **(b)** *Amd1* mRNA and AMD1 protein expression in parental and chemoresistant 4T1 and MC38 cells. RT–qPCR, n = 3 biological replicates; two-tailed unpaired Student’s t test. Immunoblots are representative of three independent experiments. **(c)** MTA concentrations in TCM from control and *Amd1*-knockdown MC38R and 4T1R cells. n = 3 biological replicates; two-tailed unpaired Student’s t test. **(d)** Schematic of the experimental design. **(e)** Representative flow cytometry plots and quantification of IFN-γ, GZMB, and PRF1 expression in CD8^+^T cells following co-culture with PMN-MDSCs generated under the indicated TCM conditions. n = 3 biological replicates/group; two-way ANOVA followed by Sidak’s multiple-comparisons test. **(f)** Representative images and quantification of primary mammosphere formation following co-culture with PMN-MDSCs generated under the indicated conditions. n = 3 biological replicates/group; one-way ANOVA followed by Sidak’s multiple-comparisons test. Scale bars, 100 μm.

To further investigate the role of AMD1-driven MTA in PMN-MDSC activity, we induced PMN-MDSCs using TCM from parental cells, resistant cells, *Amd1*-knockdown resistant cells or *Amd1*-knockdown resistant cells supplemented with MTA, followed by co-culture with CD8^+^T cells or tumor cells (Fig. 5d). Compared with resistant-cell TCM, *Amd1*-knockdown TCM generated less immunosuppressive PMN-MDSCs, allowing greater IFN-γ, Granzyme B and Perforin expression in co-cultured CD8^+^T cells and reducing tumor-sphere formation (Fig. 5e-f). However, adding MTA back to the *Amd1*-knockdown TCM restored PMN-MDSC immunosuppressive activity, as evidenced by reducing T cell effector function and increasing tumor-sphere formation (Fig. 5e-f). All these results suggest that AMD1-driven MTA production in the chemoresistant tumor cells is critical for the immunosuppressive activity of PMN-MDSCs.

### Genetic or pharmacological inhibition of AMD1 re-sensitizes the resistant tumors to chemotherapy and immunotherapy

We next tested whether suppressing tumor cell AMD1 could reverse treatment resistance in *vivo*. Resistant control and *Amd1*-knockdown tumor cells were implanted into syngeneic mice and treated with the corresponding chemotherapeutic drugs. Notably, *Amd1* knockdown restored chemotherapy sensitivity, resulting in substantially reduced tumor growth compared with resistant tumor controls (Fig. 6a and Extended Data Fig. 6a). PMN-MDSCs isolated from *Amd1*-knockdown resistant tumors were less immunosuppressive, as indicated by increased expression of cytotoxic molecules in co-cultured CD8^+^T cells and reduced tumor-sphere formation in co-cultured tumor cells, while these effects were further enhanced when *Amd1* knockdown tumors was treated with chemotherapy (Fig. 6b-c and Extended Data Fig. 6b). We then evaluated the therapeutical potential of pharmacological AMD1 blockade using CGP 48664A. Likewise, combining CGP 48664A with chemotherapy effectively blunted the growth of resistant tumors (Fig. 6d and Extended Data Fig. 6c). Moreover, the combination therapy greatly reduced intratumoral PMN-MDSC abundance and improved the cytotoxic function of tumor-infiltrating CD8^+^T cells (Fig. 6e and Extended Data Fig. 6d-e). Consistently, PMN-MDSCs isolated from combination therapy-treated tumors displayed weaker immunosuppressive activity ex vivo, allowing much higher expression of T cell effector molecules and leading to less tumor-sphere formation than PMN-MDSCs from control or chemotherapy-only tumors (Fig. 6f-g and Extended Data Fig. 6f).

**Fig. 6.**
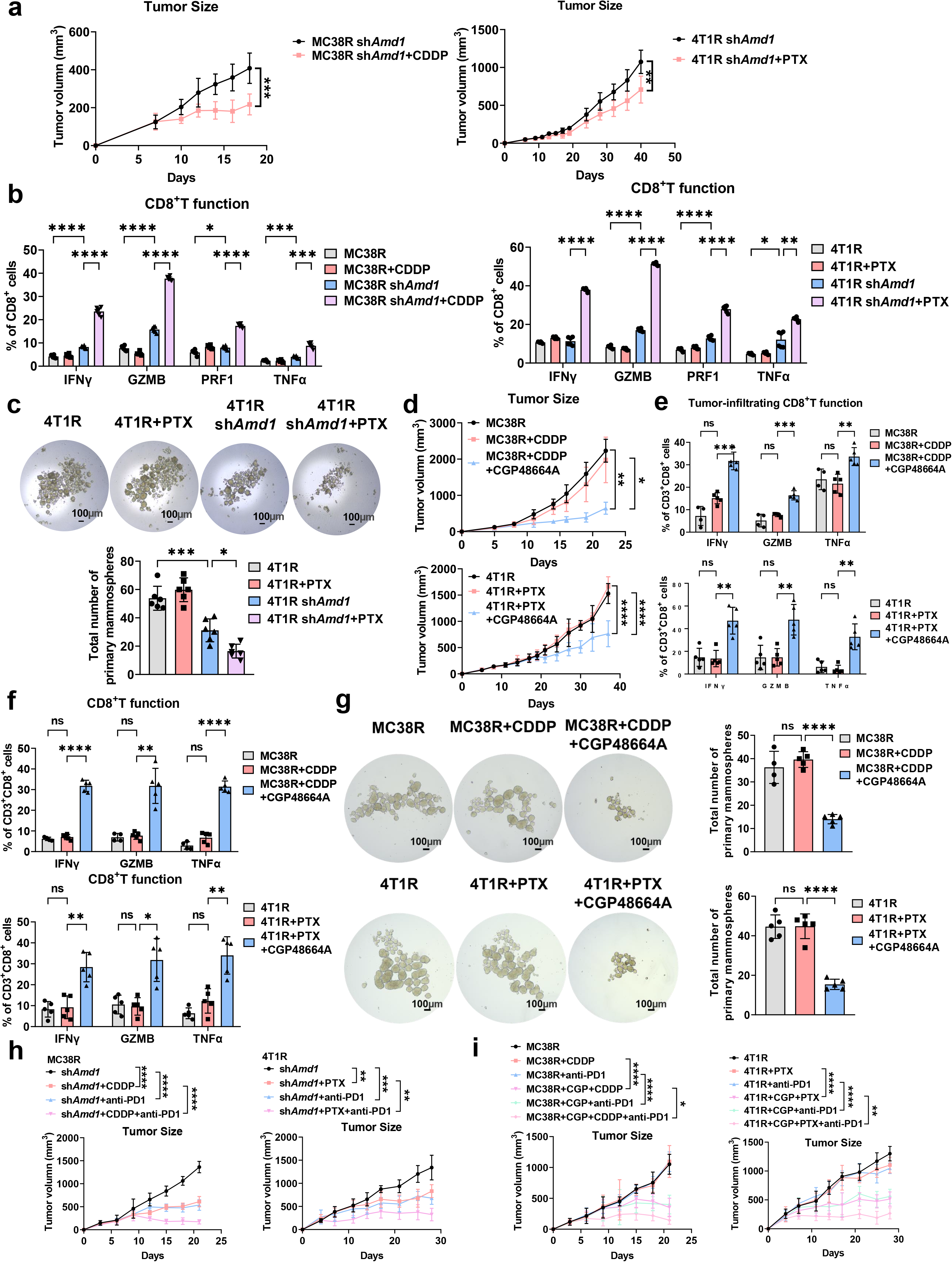
AMD1 inhibition restores sensitivity of chemoresistant tumors to chemotherapy and immunotherapy. **(a)** Tumor growth of *Amd1*-knockdown MC38R and 4T1R tumors with or without chemotherapy. n = 5-6 mice/group; repeated-measures two-way ANOVA followed by Sidak’s multiple-comparisons test. **(b)** Cytotoxic effector molecule expression in CD8^+^T cells following co-culture with PMN-MDSCs isolated from the indicated tumors. n = 5-6 mice/group; two-way ANOVA followed by Sidak’s multiple-comparisons test. **(c)** Representative images and quantification of mammosphere formation following co-culture with PMN-MDSCs isolated from the indicated 4T1R groups. n = 6 mice/group; one-way ANOVA followed by Sidak’s multiple-comparisons test. Scale bars, 100 μm. **(d)** Tumor growth of MC38R and 4T1R tumors treated with chemotherapy alone or combined with CGP48664A. n = 4-5 mice/group; repeated-measures two-way ANOVA followed by Sidak’s multiple-comparisons test. **(e)** IFN-γ, GZMB, and TNF-α expression in tumor-infiltrating CD8^+^T cells of the indicated tumors. n =4-5 mice/group; two-way ANOVA followed by Sidak’s multiple-comparisons test. **(f)** Cytotoxic effector molecule expression in CD8^+^T cells following ex vivo co-culture with PMN-MDSCs isolated from the indicated tumors. n =4-5 mice/group; two-way ANOVA followed by Sidak’s multiple-comparisons test. **(g)** Representative images and quantification of mammosphere formation following co-culture with PMN-MDSCs isolated from the indicated tumors. n =4-5 mice/group; one-way ANOVA followed by Sidak’s multiple-comparisons test. Scale bars, 100 μm. **(h)** Tumor growth of *Amd1*-knockdown MC38R and 4T1R tumors receiving chemotherapy and/or anti-PD-1. n = 5 mice/group; repeated-measures two-way ANOVA followed by Sidak’s multiple-comparisons test. **(i)** Tumor growth of MC38R and 4T1R tumors receiving chemotherapy, CGP48664A, anti-PD-1, or the indicated combinations. n = 5 mice/group; repeated-measures two-way ANOVA followed by Sidak’s multiple-comparisons test.

Given that resistant tumors were also refractory to immune checkpoint blockade, we examined whether AMD1 inhibition could restore sensitivity to anti-PD-1 therapy and chemo-immunotherapy. Strikingly, in mice bearing *Amd1*-knockdown resistant tumors, both chemotherapy and anti-PD-1 treatment monotherapy significantly inhibited tumor growth, while the combined chemo-immunotherapy achieved more pronounced tumor control (Fig. 6h and Extended Data Fig. 6g-h). These therapeutic responses were accompanied by lower PMN-MDSC abundance and improved cytotoxic function of tumor-infiltrating CD8^+^T cell (Extended Data Fig. 7a-b). Pharmacological inhibition of AMD1 similarly restored tumor responsiveness to anti-PD-1 monotherapy and chemo-immunotherapy (Fig. 6i and Extended Data Fig. 6i-j). Notably, the combination of AMD1 inhibitor, chemotherapy and anti-PD-1 therapy resulted in the strongest reduction in PMN-MDSCs and the greatest improvement of CD8^+^T cell cytotoxic function (Extended Data Fig. 7c-d). Collectively, these findings establish that genetic or pharmacological inhibition of AMD1 dismantles the immunosuppressive PMN-MDSCs, thereby re-sensitizing the resistant tumors to chemotherapy and immunotherapy.

### The AMD1–MTA–neutrophil axis confers chemoresistance in patient-derived tumor organoids

To extend our findings to human models, we established breast cancer and colorectal cancer patient-derived tumor organoids (PDOs), and analyzed their IC50 values in response to paclitaxel and oxaliplatin (Extended Data Fig. 8a), as these two agents are the most commonly used chemotherapeutic drugs in the clinical treatment for breast cancer and colorectal cancer respectively. Based on the IC50 values of these organoids, we classified the organoids into chemo-sensitive organoids (lowest IC50) and chemo-resistant organoids (highest IC50). To examine the effects of PMN-MDSCs on the tumor organoids, we isolated neutrophils (CD11b^+^ CD14^-^CD15^+^) from peripheral blood of cancer patients and treated them with vehicle or MTA. The neutrophils were then co-cultured with chemo-sensitive organoids before chemotherapeutic drug exposure (Fig. 7a). Of note, co-culture with neutrophils markedly expanded organoid number and morphology. This promoting effect was further amplified when co-cultured with MTA-treated neutrophils (Fig. 7b). To assess the organoid sensitivity across different drug concentrations in the co-culture system, the organoids were engineered to express a mCherry-luciferase reporter and co-cultured with vehicle- or MTA-treated neutrophils (Extended Data Fig. 8b). We found that chemotherapy markedly destroyed sensitive organoids cultured alone, whereas neutrophil co-culture, particularly with MTA-treated neutrophils preserved organoid number and maintained higher luciferase activity across increasing drug concentrations (Fig. 7c and Extended Data Fig. 8c). These data suggest that MTA-treated neutrophils promoted chemoresistance of tumor organoids.

**Fig. 7.**
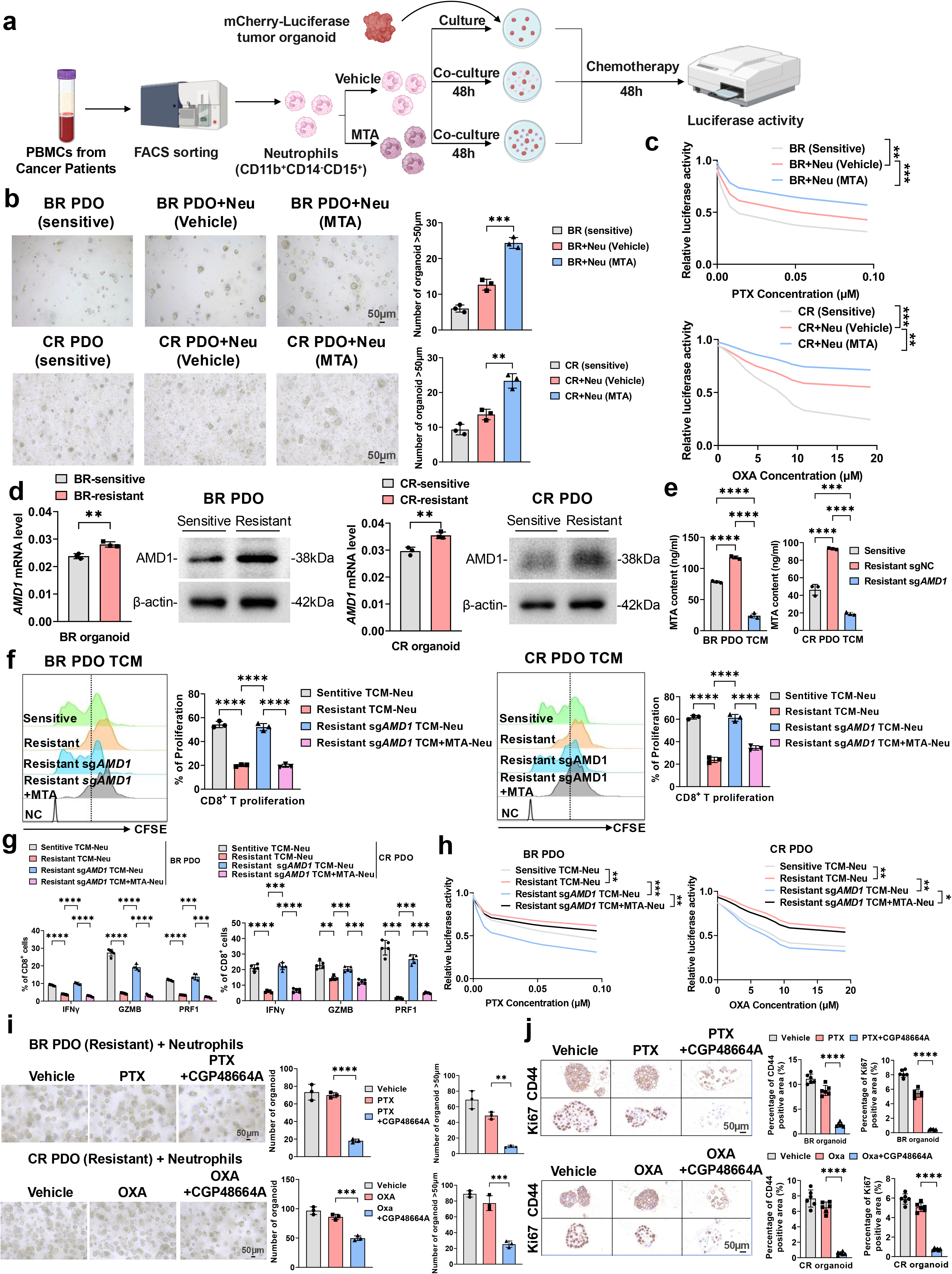
The AMD1–MTA–neutrophil axis confers chemoresistance in patient-derived tumor organoids. **(a)** Schematic illustration for the patient-derived neutrophil and PDO co-culture assay. **(b)** Representative bright-field images and quantification of PDO growth following co-culture with vehicle-or MTA-treated neutrophils. n = 3 biological replicates/group; one-way ANOVA followed by Sidak’s multiple-comparisons test. Scale bars, 50 μm. **(c)** Chemotherapy-response curves of mCherry–firefly luciferase-labelled PDOs cultured alone or with vehicle-or MTA-treated neutrophils. n = 3 biological replicates/group; two-way ANOVA followed by Sidak’s multiple-comparisons test. **(d)** *AMD1* mRNA and AMD1 protein expression in chemosensitive and chemoresistant PDOs. RT–qPCR, n = 3 biological replicates/group; two-tailed unpaired Student’s t test; immunoblots are representative of three independent experiments. **(e)** MTA concentrations in conditioned media from chemosensitive, chemoresistant, and AMD1-deficient chemoresistant PDOs. n = 3 biological replicates/group; one-way ANOVA followed by Sidak’s multiple-comparisons test. **(f)** Representative CFSE profiles and quantification of human CD8^+^T cell proliferation following co-culture with the indicated PDO-conditioned neutrophils. n = 3 biological replicates/group; one-way ANOVA followed by Sidak’s multiple-comparisons test. **(g)** IFN-γ, GZMB, and PRF1 expression in human CD8^+^T cells following co-culture with the indicated PDO-conditioned neutrophils. n = 3 biological replicates/group; two-way ANOVA followed by Sidak’s multiple-comparisons test. **(h)** Chemotherapy-response curves of drug-sensitive PDOs following co-culture with the indicated PDO-conditioned neutrophils. n = 3 biological replicates/group; two-way ANOVA followed by Sidak’s multiple-comparisons test. **(i)** Representative bright-field images and quantification of chemoresistant PDO growth co-cultured with patient-derived neutrophils following vehicle, chemotherapy, or chemotherapy plus CGP48664A treatment. n = 3 biological replicates/group; one-way ANOVA followed by Sidak’s multiple-comparisons test. Scale bars, 50 μm. **(j)** Representative Ki67 and CD44 IHC staining of chemoresistant PDOs co-cultured with patient-derived neutrophils following the indicated treatments, with quantification of positive area. n = 6 biological samples/group; six microscopic fields were analyzed for each sample and averaged to generate one value per biological sample. Data are shown as mean ± SEM; one-way ANOVA followed by Sidak’s multiple-comparisons test. Scale bars, 50 μm.

We next compared AMD1 expression between chemo-sensitive and chemo-resistant organoids. Intriguingly, both chemo-resistant breast tumor and colorectal tumor organoids exhibited higher AMD1 expression than their sensitive counterparts (Fig. 7d), along with increased MTA levels in the culture supernatant. Importantly, CRISPR-mediated AMD1 deletion in chemo-resistant tumor organoids resulted in significant reduction of MTA levels (Fig. 7e and Extended Data Fig. 8d). Moreover, neutrophils exposed to conditioned media from chemo-resistant tumor organoids potently suppressed human CD8^+^T cell proliferation and cytotoxic effector function. This suppressive activity was abrogated when the conditioned media was obtained from AMD1-deficient resistant organoids, but was recovered by supplemented MTA to the AMD1-deficient tumor organoid-conditioned media (Fig. 7f-g and Extended Data Fig. 8e). The same conditioned neutrophils were then co-cultured with luciferase-expressed sensitive organoids. Consistently, neutrophils conditioned by resistant-organoid supernatant protected sensitive organoids from chemotherapy. However, neutrophils conditioned by AMD1-deficient resistant-organoid supernatant impaired the enhanced chemoresistance, whereas MTA supplement restored the drug resistance (Fig. 7h and Extended Data Fig. 9a). These results recapitulated the tumor cell-neutrophil interactions observed in mouse models, further supporting AMD1-dependent MTA production as a key metabolic link that promotes chemoresistance by enhancing the immunosuppressive activity of PMN-MDSCs.

We next evaluated the effect of AMD1 inhibitor on the growth of tumor organoids upon chemotherapeutic drug exposure. In the resistant organoids co-cultured with neutrophils, chemotherapy alone did not impact organoid growth. Obviously, combining CGP48664A with chemotherapy prominently reduced organoid number and size. Ki67 and CD44 immunohistochemical staining further demonstrated that the growth of these organoids was severely impaired (Fig. 7i-j). Finally, analysis of scRNA-seq data verified that AMD1 expression was significantly upregulated in the tumor cells of chemoimmunotherapy-resistant patients when compared to those from sensitive patients (Extended Data Fig. 9b). Moreover, clinical datasets revealed that high AMD1 expression was associated with poor outcomes, and positively correlated with PMN-MDSC gene signatures in multiple tumor types (Extended Data Fig. 9c-d). Taken together, these findings from patient-derived tumor organoids and clinical samples extended the AMD1–MTA–PMN-MDSC axis beyond mouse models and established this pathway as a potential therapeutic vulnerability in chemoresistant tumors.

## Discussion

Chemotherapy remains an important modality in the clinical treatment of cancer. However, chemoresistance continues to pose a major challenge in clinical treatment. Under sustained selective pressure exerted by chemotherapy, tumor cells can undergo adaptive evolution through mechanisms such as the acquisition of new genetic mutations, metabolic reprogramming, increased drug efflux, and enhanced DNA damage repair^31,32^. Moreover, tumor cells that survive under chemotherapy are likely to remodel the surrounding microenvironment, thereby creating conditions that support their long-term survival within the tumor microenvironment. Here we demonstrate that chemotherapy drives metabolic reprogramming in tumor cells, leading to aberrantly elevated expression of the metabolic enzyme AMD1. This, in turn, results in excessive production and release of MTA. MTA directly binds to PTPN2 in PMN-MDSCs which are abundantly accumulated in the chemoresistant tumors, and blocks its phosphatase activity, thereby enhancing activation of JAK-STAT3 signaling. This promotes the immunosuppressive function of PMN-MDSCs, enabling tumor cells to evade T cell-mediated cytotoxicity while simultaneously enhancing tumor cell stemness to sustain their chemoresistance. Blocking AMD1-driven MTA production efficiently re-sensitizes the resistant tumors to chemotherapy and immunotherapy (Extended Data Fig. 9e).

MTA is generated principally as a coproduct of polyamine synthesis. AMD1 converts S-adenosylmethionine into decarboxylated S-adenosylmethionine, which serves as the aminopropyl donor. The transfer of the aminopropyl group to putrescine or spermidine generates spermidine or spermine, respectively, with MTA produced as a stoichiometric byproduct^18^. AMD1 therefore does not synthesize MTA directly, but controls flux through MTA-producing reactions. Interestingly, despite the marked upregulation of AMD1 in chemoresistant tumor cells, we did not detect a corresponding increase in extracellular spermine or spermidine based on the metabolomics data. Polyamines are predominantly retained within the intracellular compartment rather than freely diffusing across the plasma membrane. For instance, spermine can be converted back to spermidine either through direct oxidation by spermine oxidase (SMOX) or through an acetylation–oxidation pathway^33,34^. The absence of extracellular spermine and spermidine in our samples suggests that these polyamines are efficiently retained, metabolized, or otherwise regulated intracellularly, whereas MTA may have a distinct fate that favors extracellular accumulation. Thus, AMD1-driven polyamine metabolic reprogramming selectively generates an extracellular MTA signal through which chemotherapy-adapted tumor cells communicate with PMN-MDSCs.

The precise mechanism responsible for AMD1 upregulation in chemoresistant cells remains to be elucidated. Several mechanisms may potentially contribute to this metabolic adaptation. AMD1 is a transcriptional target of the oncogenic transcription factor MYC^35,36^. Sustained chemotherapy-induced selection for tumor cells with increased MYC activity may enhance AMD1 transcription. In addition, mTORC1 signaling has been shown to regulate AMD1 protein stability^19^, indicating that oncogenic signaling may increase AMD1 abundance at the post-translational level. Chemotherapy-induced metabolic remodeling may increase the demand for polyamines to support tumor cell survival, proliferation and stress-adaption^37–39^, which may consequently favor increased AMD1 expression. Further investigation is warranted to determine whether AMD1 upregulation is driven primarily by transcriptional activation, enhanced protein stability, or a combination of these mechanisms during the acquisition and maintenance of chemoresistance.

MTA abundance also depends on MTAP, which consumes MTA in the methionine-salvage pathway. MTAP lies adjacent to CDKN2A/B at 9p21.3 and is frequently co-deleted with these tumor suppressors^40^. In MTAP-deficient tumors, impaired MTA catabolism results in MTA accumulation, which can exert immunosuppressive effects, including inhibition of PRMT5 activity and suppression of T cell function^23,41,42^. However, this canonical mechanism does not appear to account for the MTA accumulation observed in our chemoresistant models. Notably, MTAP expression was preserved across the tumor cells examined in our study, with no evidence of MTAP loss or substantial reduction in MTAP expression. Thus, our results identify an alternative route to MTA accumulation in chemoresistant tumors. Rather than reflecting a defect in MTA catabolism caused by MTAP deficiency, the elevated extracellular MTA is more consistent with increased upstream production associated with AMD1-driven metabolic reprogramming. This finding expands the biological significance of MTA-directed immunomodulation beyond the ∼10% of MTAP-deleted cancers to the broader population of tumors that acquire chemoresistance.

Extracellular MTA can affect several immune compartments. MTA directly inhibits T cell proliferation, activation and effector function through impaired AKT signaling and altered protein methylation^43^. High MTA has also been linked to CD8^+^T cell exhaustion and chromatin closure in hepatocellular carcinoma^44^. In MTAP-deficient tumors, MTA depletion restores T cell infiltration and checkpoint-blockade sensitivity^23^. MTA additionally suppresses natural killer cells, dendritic cells and inflammatory macrophage activation^45–47^. In this study, we found that PMN-MDSCs represent the predominant immunosuppressive cell population enriched in chemoresistant tumors. Given their numerical dominance within the tumor microenvironment, we speculate that MTA may preferentially exert its immunomodulatory effects on PMN-MDSCs, thereby enhancing their suppressive activity and contributing to the maintenance of chemoresistance. However, we cannot exclude the possibility that MTA also regulates other immune or stromal cell populations. Such effects may act independently or cooperatively with PMN-MDSCs to shape an immunosuppressive microenvironment and promote chemoresistance. Further studies will be required to determine the contribution of different MTA-responsive cell populations to chemoresistance.

PMN-MDSCs are a heterogeneous population of pathologically activated neutrophils generated during cancer-associated granulopoiesis. PMN-MDSCs suppress antitumor immunity through ARG1, ROS, and S100A8/A9-dependent mediators^48,49^. The suppressive state of PMN-MDSCs is shaped by diverse soluble signals in the tumor microenvironment. For example, tumor-exosomal HSP72 can activate MDSCs through TLR2–MyD88–IL-6–STAT3 signaling and tumor-derived lactate increases PMN-MDSC recruitment^50,51^. More PMN-MDSC-specific evidence links lipid metabolism to suppressive function. GM-CSF–STAT5 signaling induces fatty acid transport protein 2 (FATP2), enabling arachidonic acid uptake and prostaglandin E2 (PGE2) synthesis required for PMN-MDSC activity^52^. PGE2 also promotes MDSC differentiation and suppression through E prostanoid (EP) receptors^53,54^. Here our findings identify MTA as a new metabolite mediating the immunosuppressive function of PMN-MDSCs. Mechanistically, MTA binds to the WPD loop of PTPN2 and disrupts its phosphatase activity, thereby facilitating JAK-STAT3 signaling. Sustained STAT3 activity drives ARG1, NOX2 and other suppressive MDSC mediators^55,56^. PTPN2 restrains cytokine signaling by dephosphorylating JAK and STAT family proteins^57^. Although PTPN2 inhibitors have been shown to potentiate CD8^+^T anti-tumor immunity and are under evaluation in the clinical trial^58,59^. Our findings raise the possibility that PTPN2 inhibition may simultaneously enhance the immunosuppressive function of myeloid cells. The dual effects on both lymphoid and myeloid compartments should be considered when developing PTPN2-targeted therapies. Beyond targeting PTPN2, MTA may regulate PMN-MDSC function through additional mechanisms, including altered methylation, purinergic signaling and metabolic reprogramming. MTA can compete with SAM at PRMT5 and reduce symmetric arginine demethylation, exerting an epigenetic regulatory role^22,40^. MTA may also act through adenosine receptors, which have been implicated in its effects on macrophages and T cells. These mechanisms may converge on the established PMN-MDSC activity^23,60^.

In summary, our study highlights that chemotherapy resistance is not solely a tumor-cell-intrinsic phenotype driven by genetic or metabolic alterations, but can also be actively maintained through metabolite-mediated communication between tumor cells and the immune microenvironment. Targeting the AMD1–MTA–PMN-MDSC axis may disrupt this self-reinforcing resistant ecosystem and provide a strategy to restore therapeutic sensitivity in refractory tumors.

## Methods

### Mice

All mice used in our experiments were six-to eight-week old, and housed under specific pathogen free (SPF) conditions in the animal facility at Ren Ji Hospital. WT C57BL/6J mice, BALB/c mice and OT-1 mice were purchased from Shanghai Model Organisms Center. All the animal studies were conducted in accordance of the protocols approved by the Animal Care and Use Committee at Ren Ji Hospital, Shanghai Jiao Tong University School of Medicine.

### Human samples

Patient derived organoids were established from the tumor tissues of breast cancer and colorectal cancer patients. Human CD8⁺T cells were purchased from Hycells Biotechnology Co., Ltd. Human peripheral blood for neutrophil isolation was obtained from healthy adult donors, breast cancer and colorectal cancer patients. All procedures involving human samples were approved by the Ethics Committee of Ren Ji Hospital, Shanghai Jiao Tong University School of Medicine (Approve NO. KY2024-126-B). The written informed consent was obtained from all participants where applicable.

### Cell lines and plasmids

4T1 mammary adenocarcinoma cell line were gifts from the laboratory of Dr. Qiujing Yu (Sichuan Provincial People’s Hospital, School of Medicine, University of Electronic Science and Technology of China). MC38/CT26 murine colorectal cell line, 293T cell line were kindly given by Dr. Bin Ma (School of Biomedical Engineering and Med-X Research Institute, Shanghai Jiao Tong University). 4T1 and CT26 cells were cultured in RPMI-1640 (Gibco), and MC38, 293T cells were cultured in DMEM (Gibco). All cells were maintained in the media supplemented with 10% fetal bovine serum (FBS, Gibco) and 100 U/ml penicillin/streptomycin (Gibco) at 37°C under 5% CO_2_ incubator. All of the cell lines were regularly tested for Mycoplasma free. Paclitaxel-resistant 4T1 (4T1R-PTX), cisplatin-resistant MC38 and CT26 (MC38R-CDDP/CT26-CDDP) derivatives were generated by continuous low-dose exposure to the corresponding drug for around 6 months. Cells were maintained in 0.014 μM paclitaxel or 0.04 μM cisplatin during the establishment of resistant derivatives. Resistance was confirmed by a rightward shift in the drug-response curve and an increased half-maximal inhibitory concentration (IC50) relative to the parental line.

Stable AMD1-knockdown 4T1R-PTX and MC38R-CDDP cells were generated by lentiviral infection. HEK293T cells were co-transfected with the pLKO.1-sh*Amd1* lentiviral plasmid, psPAX2 and pMD2.G using polyethyleneimine (Yeasen, cat#40820ES04). Lentiviral supernatants were collected at 48 and 72 hr after transfection, followed by centrifuging to remove cell debris and filtered through a 0.45μm filter. Resistant tumor cells were infected with the lentiviral supernatants in the presence of 10 μg/mL polybrene and subsequently selected with 10 μg/mL puromycin to establish stable AMD1-knockdown cell lines. Knockdown efficiency was confirmed by RT–qPCR and immunoblotting. The shRNA sequence was listed in Table S1.

An EGFP-tagged mouse PTPN2 overexpression plasmid (pLV-CMV-EGFP-m*Ptpn2*) and the corresponding control vector were constructed for transient expression in PMN-MDSCs. The PTPN2 overexpression plasmid was introduced into PMN-MDSCs by electroporation as described below.

### Cell electroporation

For PTPN2 overexpression, 2 × 10⁶ PMN-MDSCs were electroporated with 4 μg PTPN2 overexpression plasmid using a Gene Pulser Xcell electroporation system (Bio-Rad). Plasmid DNA was mixed with EZ Trans transfection reagent (Liji Biotech, cat#AC04L092) at a ratio of 1 μg DNA to 1 μL reagent before electroporation. The cell– DNA mixture was transferred into a 2 mm electroporation cuvette and electroporated at 180 V and 500 μF with infinite resistance. Following electroporation, cells were immediately transferred to complete culture medium and cultured for 48 hr before subsequent experiments.

### siRNA transfection

siRNA was synthesized by Shanghai BiOligo Biotech Co., Ltd. The sequences were listed in Table S1. Cells were seeded in 6-well plates and transfected with siRNA at a final concentration of 15 nM using Lipofectamine™ RNAiMAX Transfection Reagent (Thermo Fisher Scientific, cat#13778030) according to the manufacturer’s instructions. Cells were collected 48 hr after transfection for subsequent experiments.

### Tumor conditioned medium preparation

Tumor-conditioned medium (TCM) was prepared from tumor cells cultured in 150 mm dishes containing 20 ml of complete medium. At approximately 80% confluence, the cells were washed three times with phosphate-buffered saline and cultured in serum-and antibiotic-free basal medium for 48 h. The conditioned medium was collected and filtered with a 0.22 μm filter and then stored at -80℃ until use.

### Syngeneic tumor models and treatments

For syngeneic transplanted breast tumor models, 5x10⁵ parental 4T1, 4T1R-PTX or 4T1R-PTX sh*Amd1* cells suspended in 100 μL of PBS were injected into the fourth mammary fat pad of female BALB/c mice. For syngeneic colorectal tumor models, 5x10⁵ parental MC38, MC38R-CDDP or MC38R-CDDP sh*Amd1* cells were subcutaneously injected into male C57BL/6J mice. For the CT26R colorectal tumor model, 5 × 10⁵ CT26R-CDDP cells were subcutaneously injected into male BALB/c mice. For chemotherapy treatment, mice with palpable tumors were injected intraperitoneally (i.p.) with 7 mg/kg paclitaxel (MedChemExpress, cat#HY-B0015) or 7 mg/kg cisplatin (MedChemExpress, cat#HY-17394) every other day. For the AMD1 inhibition, CGP 48664A (MedChemExpress, cat#HY-13746B) was dissolved in ddH_2_O and intraperitoneally administered at 20 mg/kg every other day. For the Ly6G⁺ cell depletion in *vivo*, mice were injected intraperitoneally with 200 μg/mouse anti-Ly6G antibody (Bio X Cell, cat#BE0075) or the isotype control (Bio X Cell, cat#BE0089) one day before tumor-cell inoculation and every other day thereafter. Anti-PD-1 mAb (Bio X Cell, cat#BE0146) was intraperitoneally administrated with 150 μg/mouse every four days. For MTA-supplementation in *vivo*, mice with palpable tumors were injected intraperitoneally with MTA (MedChemExpress, cat#HY-16938) at 30 mg/kg twice a week in a vehicle consisting of 10% DMSO, 40% PEG300, 5% Tween-80 and 45% saline. Tumor volumes were measured along largest diameter and perpendicular diameter (a, b) and calculated as V =axbxb/2. The mice were euthanized and the tumor tissues were collected until the tumor volume reached ∼1000mm^3^.

### Flow cytometry analysis and sorting

Following mouse euthanasia, tumors were immediately dissected. Tumors were finely minced and incubated in DMEM (Gibco) supplemented with collagenase IV (Sigma, cat#C5138) at 1.5 mg/mL, and DNase I (Roche, cat#11284932001) at 100 µg/mL on a shaker at 250 rpm, 37℃ for 45 minutes. Complete medium with FBS was added to the mixture to terminate digestion, and the mixture was filtered with a 70 µm cell strainer, washed with PBS once, and resuspended with 1% BSA-PBS containing 1 mM EDTA. The isolated cells were incubated with anti-mouse CD16/32 antibody for 15 min on ice to block nonspecific Fc receptor binding. Cells were then stained with cell surface markers: anti-CD45-BV510 (Biolegend, Clone30-F11, cat#103137), anti-CD45-FITC (Biolegend, Clone30-F11, cat#103108), anti-CD3-AF700 (Biolegend, Clone17A2, cat#100216), anti-CD4-APC (Biolegend, CloneGK1.5, cat#100412), anti-CD8-PE (Biolegend, Clone YTS156.7.7, cat#126608), anti-CD4-PerCP/Cyanine5.5 (Biolegend, Clone CloneGK1.5, cat#100434), anti-CD11b-APC (Biolegend, CloneM1/70, cat#101212), anti-Ly-6G-AF700 (Biolegend, Clone1A8, cat#127622), anti-Ly-6C-PE (Biolegend, CloneHK1.4, cat#128008), anti-F4/80-PerCP/Cyanine5.5 (Biolegend, Clone BM8, cat#123126), anti-CD45R/B220-PerCP (Biolegend, CloneRA3-6B2, cat#103233), and incubated on ice for 30 min. After washing with PBS, cells were resuspended with FACS buffer containing viability dye DAPI (1:2000) for 10 min and acquired by LSR Fortessa flow cytometry (BD Biosciences). Data were analyzed with Flowjo software (Tree Star).

For intracellular staining of T cell cytotoxic molecules, 5x10^5^ cells were pre-incubated with Cell Activation Cocktail (Biolegend, cat#423303) for 8 hr, and stained with surface markers anti-CD45-BV510, anti-CD3-AF700 and anti-CD8-PE antibodies for 30 min. The cells were washed with PBS and fixed with fixation buffer (eBioscience, cat#00-8222-49) for 30 min. After that, cells were stained with anti-Granzyme B-FITC (Biolegend, CloneQA18A28, cat#396404), anti-IFN-γ-APC (Biolegend, CloneXMG1.2, cat#505809), anti-TNF-α-BV605 (Biolegend, CloneMP6-XT22, cat#506329) and anti-Perforin-PB (Biolegend, CloneS16009B, cat#154408) antibodies diluted in permeabilization buffer for 30 min. Cells were washed with PBS and acquired by LSR Fortessa flow cytometry.

For tumor cell staining, anti-CD133-APC (Biolegend, Clone315-2C11, cat#141207) and anti-CD44-FITC (Biolegend, CloneNIM-R8, cat#156007) antibodies were used.

For sorting of intratumoral PMN-MDSCs, anti-CD45-FITC (Biolegend, Clone30-F11, cat#103108), anti-CD11b-APC (Biolegend, CloneM1/70, cat#101212), anti-Ly-6G-PerCP/Cy5.5 (Biolegend, Clone1A8, cat#127615) and anti-Ly-6C-PE (Biolegend, CloneHK1.4, cat#128008) antibodies were used.

For sorting of human neutrophils from PBMCs, anti-CD11b-RB705 (BD Pharmingen, CloneD12, cat#757194), anti-CD14-RY586 (BD Pharmingen, Clone MφP9, cat#753381) and anti-CD15-FITC (BD Pharmingen, CloneHI98, cat#555401) antibodies were used. CD11b⁺CD14⁻CD15⁺ cells were sorted.

### Mouse neutrophil isolation and PMN-MDSC induction

Neutrophils were isolated from the bone marrow of wild-type C57BL/6J or BALB/c mice using the EasySep Mouse Neutrophil Enrichment Kit (STEMCELL, cat#19762A). To induce PMN-MDSCs, purified neutrophils were cultured for 3 days in a 1:1 mixture of complete RPMI 1640 medium and TCM, supplemented with 20 ng/ml granulocyte– macrophage colony-stimulating factor (GM-CSF). For the other treatments, freshly isolated neutrophils or TCM-induced PMN-MDSCs were treated with 250 μM MTA for 24 h. To inhibit STAT3 signaling, cells were pretreated with 5 μM Stattic (MedChemExpress, cat#HY-13818) for 4 h before MTA exposure.

### CD8^+^T cell isolation and co-culture experiments

OT-1 CD8^+^T cells were isolated from the spleens of OT-1 mice using the EasySep Mouse CD8^+^T Cell Isolation Kit (STEMCELL, cat#19858). Purified OT-1 CD8^+^T cells were co-cultured with PMN-MDSCs at a T cell: PMN-MDSC ratio of 1:3 for 72 h in complete medium supplemented with 2 ng/ml IL-2 (PeproTech, cat#212-12) and 2 µg/ml OVA peptide (Ana Spec, cat#AS-60193-1). After co-culture, cells were washed, resuspended in staining buffer and stained with anti-CD8b-PE (Biolegend, CloneYTS156.7.7, cat#126608) for 30 min on ice. For proliferation assays, purified OT-1 CD8^+^T cells were labelled with CFSE (Biolegend, cat#423801) before co-culture. After 72 h, T-cell proliferation was quantified by CFSE dilution in the FITC channel. For assessment of T cell effector function, cells were fixed, permeabilized, and subsequently stained intracellularly with anti-Granzyme B-FITC, anti-IFN-γ-APC, anti-TNF-α-BV605 and anti-Perforin-PB as described above.

### Tumor sphere formation

Tumor sphere formation assays were performed as previously described^9^. Briefly, sorted or TCM-induced PMN-MDSCs were co-cultured with tumor cells at a PMN-MDSC: tumor-cell ratio of 4:1. Tumor cells were seeded at 500 cells per well in 96-well plates and cultured for 6 d in MammoCult Human Medium (STEMCELL, cat# 05620) supplemented with 4 µg/ml heparin (STEMCELL, cat#07980), 1 µg/ml hydrocortisone (MedChemExpress, cat#HY-N0583) and 1% penicillin–streptomycin. Fresh complete medium was added every 2–3 d. Spheres were imaged and counted at endpoint.

### Drug response assays

For tumor cell drug-response assays, MC38, MC38R-CDDP tumor cells were seeded at 5×10³ cells per well in 96-well plates and allowed to attach overnight. Cells were then treated with cisplatin at 0.01, 0.02, 0.05, 0.1, 0.2, 0.5, 5, 10 or 20 μM for 24 h. Similarly, 4T1, 4T1R-PTX tumor cells were seeded at 5 × 10³ cells per well, allowed to attach overnight and treated with paclitaxel at 0.01, 0.05, 0.1, 0.5, 1, 1.5, 2 or 5 μM for 24 h. Cell viability was measured using Cell Counting Kit-8 (Beyotime, cat#C0037) according to the manufacturer’s instructions. For PMN-MDSC co-culture assays, PMN-MDSCs and tumor cells were mixed at a 1:1 ratio and seeded in 96-well plates. After 24 h of co-culture, the wells were washed twice with PBS and the remaining adherent tumor cells were subjected to the corresponding cisplatin or paclitaxel drug-response assay described above, and cell viability was measured using Cell Counting Kit-8.

### Isolation of tumor interstitial fluid (TIF)

Mice tumor interstitial fluids were collected as previously described^61^. Briefly, tumors were collected and washed twice with PBS. Equal weight tissues were dissected and placed on 20 µm cell strainers fixed on top of 50 ml conical tubes and subjected to low speed centrifugation at 100 x g for 15 min at 4℃. The collected liquid was further centrifuged at 10, 000 x g for 10 min at 4℃ to remove any tissue debris. The fluid isolated from fresh tumor tissues were measured for MTA content.

### Untargeted and targeted metabolomic analyses

TCM samples were collected as described above. Untargeted metabolomic profiling was performed by Applied Protein Technology Co., Ltd. (Shanghai, China). Samples were subjected to metabolite extraction with cold methanol/acetonitrile, followed by centrifugation and vacuum drying. The dried extracts were reconstituted in acetonitrile/water and analyzed by UHPLC coupled to high-resolution mass spectrometry in both positive and negative electrospray ionization modes. Quality-control samples were prepared by pooling aliquots from individual samples and analyzed throughout the analytical sequence to monitor instrument stability. Raw MS data were converted to mzXML format using ProteoWizard MSConvert and processed using XCMS for peak detection and alignment. Isotopes and adducts were annotated using CAMERA, and metabolites were identified by matching accurate mass (mass error <10 ppm) and MS/MS spectra against an in-house database containing authentic standards. Differential metabolites were identified based on a variable importance in projection (VIP) score >1 and *P* < 0.05.

For targeted metabolomic analysis, MTA concentrations in TCM and tumor interstitial fluid (TIF) were quantified by Applied Protein Technology Co., Ltd. using a targeted LC–MS/MS assay. Liquid samples were extracted with cold methanol/acetonitrile (1:1, v/v) containing stable-isotope internal standards, centrifuged at 14,000 × g for 20 min at 4°C, and the resulting supernatants were dried under vacuum. The dried extracts were reconstituted in acetonitrile/water (1:1, v/v) and analyzed by UHPLC–MS/MS. MTA was quantified using multiple-reaction monitoring (MRM), and quantitative data were processed using MultiQuant or Analyst software.

### Quantitative reverse transcription PCR

Total RNA from tumor cells or PMN-MDSCs was extracted using Trizol (Vazyme, cat#R411-02) according to the manufacturer’s instructions and reversed transcribed with HiScript III All-in one RT SuperМix Perfect for qPCR kit (Vazyme, cat#R333-01) to make cDNA. RT-qPCR was performed using SYBR qPCR Master Mix (Vazyme, cat#Q711-03). The expression of each gene was calculated based on the cycle threshold which was set within the linear range of DNA amplification. The relative expression was calculated by the cycle threshold method, with normalization of raw data to a housekeeping gene gapdh (2^-ΔΔCt^). The primer sequences are provided in Table S1.

### Western Blotting

Cells were lysed with RIPA buffer containing protease inhibitor cocktail. The whole-cell lysates were boiled for 10 min and loaded on SDS-PAGE gels, electrophoresed, transferred onto PVDF membranes, blocked with 5% BSA-TBST or nonfat milk-TBST, and further incubated with the primary and secondary antibodies. After that, the membranes were washed with TBST for 5 times. The blots were developed using ECL reagent with Bio-Rad ChemiDoc Touch machine. The following antibodies were used: Phospho-JAK1 (1:1000, CST, cat#74129T), JAK1 (1:1000, CST, cat#3344T), Phospho-STAT3 (1:2000, CST, cat#9145), STAT3 (1:2000, CST, cat#4904), β-Actin (1:1000, Proteintech, cat#20536-1-AP), AMD1 (1:1000, Proteintech, cat#11052-1-AP), PTPN2 (1:10000, Proteintech, 11214-1-AP), MTAP (1:2000, Proteintech, 11475-1-AP), HRP Goat Anti-Rabbit IgG (H+L) (1:20000, Abclonal, cat#AS014).

### Biotin–MTA affinity purification and LC–MS/MS analysis

Biotin-labeled MTA and biotin were synthesized by Chongqing Yusi Medicine Technology. PMN-MDSCs were incubated with biotin-labeled MTA or biotin control, while untreated cells were included as an additional negative control. Cells were subsequently collected and lysed, and equal amounts of protein lysates were incubated with streptavidin magnetic beads for affinity purification. After extensive washing, bead-bound proteins were subjected to on-bead tryptic digestion and LC–MS/MS analysis at Applied Protein Technology Co., Ltd. (Shanghai, China). Peptides were separated using a nanoElute LC system and analyzed on a timsTOF Pro mass spectrometer (Bruker) operated in PASEF mode. Raw MS data were analyzed using MaxQuant (version 1.6.14), with peptide-and protein-level false discovery rates set at 1%. Candidate MTA-associated proteins were defined as proteins specifically detected in the biotin–MTA group but not in either the biotin-control or negative-control groups. PTPN2 association was further validated by streptavidin pull-down followed by immunoblotting with an anti-PTPN2 antibody.

### Molecular docking

Molecular docking was performed using AutoDock Vina. The MTA SMILES string was obtained from PubChem and converted to PDBQT format using Open Babel. The mouse PTPN2 structure was obtained from UniProt and prepared in AutoDockTools by adding polar hydrogens and assigning Gasteiger charges. PTPN2 was defined as the receptor and MTA as the ligand. A global docking strategy was used, with the grid box defined to encompass the entire PTPN2 structure. Six predicted docking poses were retained and ranked according to their Vina scores. The resulting poses were visualized in PyMOL.

### PTPN2 phosphatase activity

PTPN2 phosphatase activity was measured using p-nitrophenyl phosphate (pNPP) as a chromogenic substrate (Sigma-Aldrich, cat#20-106). The assay buffer was freshly prepared and contained 50 mM Tris–HCl (pH 8.0), 100 mM NaCl, 1 mM EDTA and 1 mM dithiothreitol. A recombinant PTPN2 stock solution (0.25 mg/mL) was added at 0.5, 1, 2, 4, 6 or 8 µL per reaction. Reactions were conducted in the absence or presence of 250 µM MTA. Reactions lacking both PTPN2 and MTA served as blanks. PTPN2 phosphatase activity was determined by measuring the absorbance generated by pNPP hydrolysis at 405 nm.

### Patient-derived organoid culture and drug response

Breast cancer and colorectal cancer patient-derived organoids (PDOs) were established and maintained as previously described^62^. Fresh surgically resected tumor tissues were minced and incubated in Tissue Digestion Solution (OneTar, cat# DB100) at 37 °C for 15-40 min. The digested tissue was passed through a 100 µm cell strainer, washed with Washing Buffer (OneTar, cat#QX100), and embedded in Matrigel (Corning, cat#356231). The cell suspension was adjusted to approximately 1.5 × 10⁶ single cells/mL, washed, and seeded in 20 µL of Matrigel per well of a 48-well plate. Breast cancer and colorectal cancer PDOs were maintained in their respective tissue-specific media (OneTar, cat#T2305-BC100 and T2301-CO100). We refreshed the medium every 2–4 days. PDOs were passaged approximately every 7 days. Briefly, organoids were incubated with TrypLE Express (Gibco, cat#12605028) for 1-3 min and mechanically dissociated by pipetting. The resulting fragments were washed at least twice to remove residual Matrigel, resuspended in fresh Matrigel, and replated. Bright-field images were acquired throughout the culture period.

For baseline chemosensitivity profiling, 5,000 organoid cells were plated per well of a 96-well plate and cultured for 3 d before drug exposure. Breast cancer organoids were exposed to different dosage of paclitaxel for 48 h; whereas colorectal cancer organoids were exposed to different dosage of oxaliplatin for 48 h. Cell viability was measured with 3D CCK-8 reagent (APExBIO, cat#K2270), PDOs were subsequently classified as sensitive or resistant according to their IC50 values.

### CRISPR–Cas9 deletion of AMD1 in patient-derived organoids

An AMD1-targeting single-guide RNA (sgRNA) was cloned into BsmBI-digested lentiCRISPR v2 using the following oligonucleotides: 5′-CACCGCGAGCGCTAGCTAATGTCGC-3′ and 5′-AAACGCGACATTAGCTAGCGCTCGC-3′. Plasmid DNA was prepared using the FastPure Plasmid Mini Kit (Vazyme, cat#DC205-01). HEK293T cells were co-transfected with the lentiCRISPR v2-sgAMD1 construct and the lentiviral packaging plasmids psPAX2 and pMD2.G. Transfection was performed using EZ Trans transfection reagent (Liji Biotech, cat#AC04L092). Viral supernatants were collected at 48 and 72 h after transfection, pooled and concentrated.

Drug-resistant PDOs were transduced overnight in a total volume of 250 µL culture medium containing 50 µL concentrated viral suspension. The organoids were subsequently replated in Matrigel and allowed to recover for 3 d. Transduced PDOs were selected with puromycin for 2 weeks. Puromycin was used at 20 µg/mL for breast cancer PDOs and 2 µg/mL for colorectal cancer PDOs. AMD1 depletion was confirmed by immunoblotting.

### mCherry-Luciferase Engineering in PDOs

Lentiviral vectors co-expressing mCherry and luciferase driven by the EF1α promoter (Lenti-EF1a-mcherry-luciferase-Puro) were used for organoid transduction. PDOs were dissociated and collected by centrifugation. The organoid pellet was resuspended in 200 μL culture medium containing 50 μL lentiviral suspension and incubated overnight at 37°C. The following day, organoids were washed with PBS, collected by centrifugation and resuspended in Matrigel for replating. Organoids were subsequently cultured and passaged routinely, and mCherry-positive organoids were sorted for downstream functional assays.

### Human neutrophil and PDO co-culture

Patient-derived neutrophils were isolated and sorted from peripheral blood as described above. For MTA-treatment experiments, purified neutrophils were incubated with vehicle or 250 µM MTA for 24 h.

For neutrophil-mediated organoid protection in response to chemotherapy, 10,000 vehicle-or MTA-pretreated neutrophils were co-cultured with 5,000 mCherry–firefly luciferase-labelled drug-sensitive organoid cells for 48 h. Chemotherapy was subsequently added, and the cultures were maintained for an additional 48 h. Breast cancer PDOs were treated with paclitaxel at 0.001, 0.005, 0.01, 0.05 or 0.1 µM. Colorectal cancer PDOs were treated with oxaliplatin at 2, 4, 8, 10 or 20 µM. Organoid morphology, number and size were quantified from bright-field images. Following organoid recovery and dissociation, firefly luciferase activity was measured under aerobic conditions using D-luciferin with ATP and Mg^2+^. Luminescence was recorded and used as a measure of viable organoid burden.

To evaluate the effects of AMD1 inhibition, drug-resistant PDOs were combined with untreated patient-derived neutrophils at a PDO ratio of 2:1. Co-cultures were divided into vehicle, chemotherapy and CGP48664A plus chemotherapy groups. In the combination group, 10 µM CGP48664A was added 48 h before chemotherapy and maintained throughout the subsequent chemotherapy exposure. Breast cancer PDOs were treated with 0.01 µM paclitaxel, whereas colorectal cancer PDOs were treated with 5 µM oxaliplatin. Bright-field images were acquired after 48 h of chemotherapy exposure, and organoid number and size were quantified. Ki67 and CD44 expression in the organoids were assessed by immunohistochemical staining.

### PDO–neutrophil and human T cell co-culture

Conditioned media were collected from breast cancer and colorectal cancer organoids representing the sensitive, resistant, sg*AMD1*-resistant, and sg*AMD1*-resistant plus MTA supplementation. Healthy donor-derived neutrophils were incubated with the corresponding conditioned medium for 48 h before co-culture with human CD8^+^T cells. Neutrophils and CD8^+^T cells were co-cultured at a ratio of 3:1, using 3 × 10^5^ neutrophils and 1 × 10^5^ T cells per condition. Before co-culture, 96-well plates were coated with 2 µg/mL anti-human CD3 antibody (Biolegend, CloneOKT3, cat#317325). After cell seeding, 2 µg/mL anti-human CD28 antibody (Biolegend, CloneCD28.2, cat#302933) and 2 ng/mL recombinant human IL-2 (PeproTech, cat#200-02) were added. Cells were co-cultured for 72 h.

For proliferation assays, CD8^+^T cells were labelled with CFSE before co-culture. After 72 h, cells were surface-stained with anti-human CD8-PE, and T-cell proliferation was quantified by CFSE dilution.

For assessment of T cell effector function, cells were surface-stained with anti-CD8-PE (Biolegend, CloneSK1, cat#344705), fixed and permeabilized. Intracellular staining was performed using anti-human Perforin-APC (Biolegend, CloneB-D48, cat#353311), anti-human IFN-γ-BV605 (Biolegend, Clone4S.B3, cat#502535), anti-human TNF-α-BV421 (Biolegend, CloneMAb11, cat#502931) and anti-human Granzyme B-FITC (Biolegend, CloneQA18A28, cat#396404).

For neutrophil-mediated organoid protection assays, 1 × 10⁴ conditioned neutrophils were co-cultured with 5 × 10³ mCherry-luciferase-labelled drug-sensitive PDO cells for 48 hr. Breast cancer PDOs were subsequently treated with paclitaxel at 0.001, 0.005, 0.01, 0.05 or 0.1 μM, whereas colorectal cancer PDOs were treated with oxaliplatin at 2, 4, 8, 10 or 20 μM for an additional 48 hr. Viable organoid burden was quantified by firefly luciferase activity.

### RNA sequencing

RNA sequencing was performed on vehicle- and MTA-treated PMN-MDSCs by LC-Bio Technology Co., Ltd. (Hangzhou, China). Total RNA was extracted using TRIzol reagent and subjected to quality assessment using a NanoDrop spectrophotometer and an Agilent 2100 Bioanalyzer. Samples with an RNA integrity number (RIN) >7.0 were used for library preparation. Poly(A) RNA was enriched using oligo(dT) magnetic beads, fragmented and reverse-transcribed into cDNA. Sequencing libraries were constructed and subjected to 150-bp paired-end sequencing on an Illumina NovaSeq platform. Raw sequencing reads were filtered using fastp to remove adaptor contamination and low-quality reads. Clean reads were aligned to the mouse reference genome using HISAT2, and transcript abundance was quantified using StringTie. Differentially expressed genes were identified using DESeq2 with |log2 fold change| ≥1 and a false discovery rate (FDR) <0.05. Gene Ontology (GO) and Kyoto Encyclopedia of Genes and Genomes (KEGG) pathway enrichment analyses were performed for differentially expressed genes. Gene set enrichment analysis (GSEA) was performed using gene sets obtained from the Molecular Signatures Database (MSigDB). Genes were ranked according to differential expression, and pathway enrichment was evaluated based on the normalized enrichment score (NES) and FDR.

### Public transcriptomic and clinical datasets

Publicly available pretreatment transcriptomic datasets with annotated responses to neoadjuvant chemotherapy were analyzed using the Cancer Treatment Response gene signature DataBase (CTR-DB 2.0). Three breast cancer cohorts, GSE162187, GSE20271 and GSE20194, were included. In GSE162187, breast tumor biopsies were obtained from patients receiving anthracycline/cyclophosphamide-based neoadjuvant chemotherapy followed by taxane treatment, including paclitaxel or docetaxel. Patients achieving a pathological complete response (pCR) were classified as responders, whereas those with residual disease were classified as non-responders. In GSE20271, patients with stage I–III breast cancer received neoadjuvant chemotherapy consisting of weekly paclitaxel followed by fluorouracil, doxorubicin or epirubicin, and cyclophosphamide (T/FAC or T/FEC), or FAC/FEC alone. In GSE20194, patients with newly diagnosed stage I–III breast cancer received approximately six months of neoadjuvant chemotherapy including paclitaxel, 5-fluorouracil, cyclophosphamide and doxorubicin. Response classifications were based on the curated clinical annotations provided by CTR-DB, with pCR categorized as response and residual disease or non-pCR categorized as non-response.

Tumor microenvironment cell abundance was estimated using the MCP-counter algorithm implemented in CTR-DB 2.0. MCP-counter-derived neutrophil and CD8⁺T cell abundance scores were compared between responders and non-responders. Differences in inferred cell abundance between the two response groups were assessed using two-sided statistical testing implemented by CTR-DB, with Benjamini– Hochberg correction for multiple comparisons. An adjusted P value < 0.05 was considered statistically significant.

### scRNA-seq data analysis

Public single-cell RNA-sequencing datasets from patients with colorectal cancer (GSE236581) and breast cancer (GSE302453) treated with chemo-immunotherapy were obtained from the GEO. In these datasets, colorectal cancer patients were received with Pembrolizumab (anti-PD-1) + Capecitabine + Oxaliplatin combination therapy, breast cancer patients were received with Pembrolizumab (anti-PD-1) + Carboplatin + Paclitaxel + Anthracycline combination therapy. Samples were classified as treatment-sensitive or treatment-resistant according to the clinical response annotations provided with the original datasets. Single-cell transcriptomic analyses were performed in R using Seurat.

Cells with 200–5,000 detected genes, fewer than 20,000 total RNA counts and less than 20% mitochondrial transcripts were retained. Data were normalized, and the top 2,000 highly variable genes were identified using the variance-stabilizing transformation method, followed by data scaling and principal component analysis (PCA). Sample-associated batch effects were corrected using Harmony. The first 30 Harmony dimensions were used for nearest-neighbor graph construction, unsupervised clustering and uniform manifold approximation and projection (UMAP), with a clustering resolution of 0.8. Cell identities were assigned based on canonical marker expression, supported by SingleR reference annotation, and manually curated. Cell-type proportions were calculated for each patient and subsequently summarized according to treatment response.

Functional states of PMN-MDSCs, T cells and malignant cells were evaluated using prespecified, dataset-specific gene signatures, and corresponding signature scores were calculated using UCell. Across the two datasets, genes used to characterize the PMN-MDSC immunosuppressive program included *SIRPA, S100A8, S100A9, LGALS9, IL10, PTGS2, NCF2, CD300LD* and *NCF1*. Genes used to characterize T-cell activation and cytotoxicity included *TNFRSF9, ICOS, IL2RA, CD69, IFNG, GZMB, GZMH, GZMA, PRF1, CCL4* and *CTSW*. Tumor stemness, adhesion and resistance-associated programs were evaluated using *SOX9, PROM1, CD44, VIM, FN1, PTK2, LDHA, KLF4, ALDH1A1, SNAI1, NDRG1* and *CDK1*. Individual gene expression and functional signature scores were compared between treatment-sensitive and treatment-resistant groups.

Cell–cell communication patterns were evaluated using CellChat-derived interaction weights to characterize intercellular signaling among the annotated cell populations. Treatment-sensitive and treatment-resistant groups were analyzed separately, and communication strengths between major cell populations were summarized at the cell-type level. Analyses focused on PMN-MDSCs and their interactions with T cells, cancer cells, endothelial cells, smooth muscle cells, monocytes/macrophages, fibroblasts and B cells. Interaction strengths were compared between treatment-sensitive and treatment-resistant groups and visualized as centered cell–cell communication networks, with edge widths representing the relative strength of intercellular communication. A common maximum interaction weight was applied to the treatment-sensitive and treatment-resistant networks to ensure directly comparable edge-width scaling between the two groups.

Individual gene expression and functional signature scores were compared between treatment-sensitive and treatment-resistant groups. In addition, normalized AMD1 expression was extracted from annotated malignant cells in both datasets and compared between treatment-sensitive and treatment-resistant groups using a two-sided Wilcoxon rank-sum test.

### Statistics and reproducibility

All described results are representative of at least three independent experiments. The number of samples (n) were described in detail for each figure panel. No statistical method was used to predetermine sample size. The experiments were not randomized, except that the mice were randomly grouped before treatment. Investigators were not blinded to allocation during the experiments and outcome assessments. Data were presented as mean±SEM. Statistical analysis was performed using GraphPad prism 9.0 software. Two-tailed unpaired Student’s t-test was used for the comparison between two groups. One-way analysis of variance (ANOVA) or two-way ANOVA followed by the Sidak’s test was used for the multiple comparisons. Repeated-measures two-way ANOVA (mixed model) followed by the Sidak’s multiple comparisons test was used for analysis of the tumor growth curve. Statistical significance was denoted with *(*P* < 0.05), **(*P* < 0.01), ***(*P* < 0.001), **** (*P*<0.0001) in the figures.

## Data availability

Raw RNA-seq data have been deposited to the Gene Expression Omnibus under accession number GSE348379. The mass spectrometry proteomics data have been deposited to the ProteomeXchange Consortium via the PRIDE partner repository with the dataset identifier PXD084643. All other data supporting the findings of this study are available from the corresponding authors upon reasonable request.

## Acknowledgements

This work was supported by funds from the National Natural Science Foundation of China, NSFC 32422029 (J.G.), 82472915 (J.G.), 82272997 (H.L.), and 824B208 (C.H.); Noncommunicable Chronic Diseases-National Science and Technology Major Project 2026ZD0557103 (J.G.); Key Laboratory of Systems Medicine for Cancer “Basic-clinical Co-innovation Team SMC-BCC-02 (J.G.); the National 111 Project of China (B21024).

## Author contributions

J.G. and D.X. conceived the study and designed the research. Y.W. and W.Q. designed the experimental approach, performed the experimental work, and analyzed the data. B.Y., G.W. and D.X. provided and coordinated the organoid culture resources and methods. C.H., H.L., H.L, R.Z., L.L., P.L., N. S., S.B., S. G. coordinated the experimental work. J.G. and Y.W. interpreted data. J.G. and Y.W. wrote the manuscript. All the authors discussed and read the manuscript.

## Declaration of Interests

The authors declare no potential conflict of interests.

## Notes

### Competing Interest Statement

The authors have declared no competing interest.

