## Supplemental figures and figure legends for "AMD1-driven MTA production sustains chemoresistance by rewiring PMN-MDSC immunosuppression"

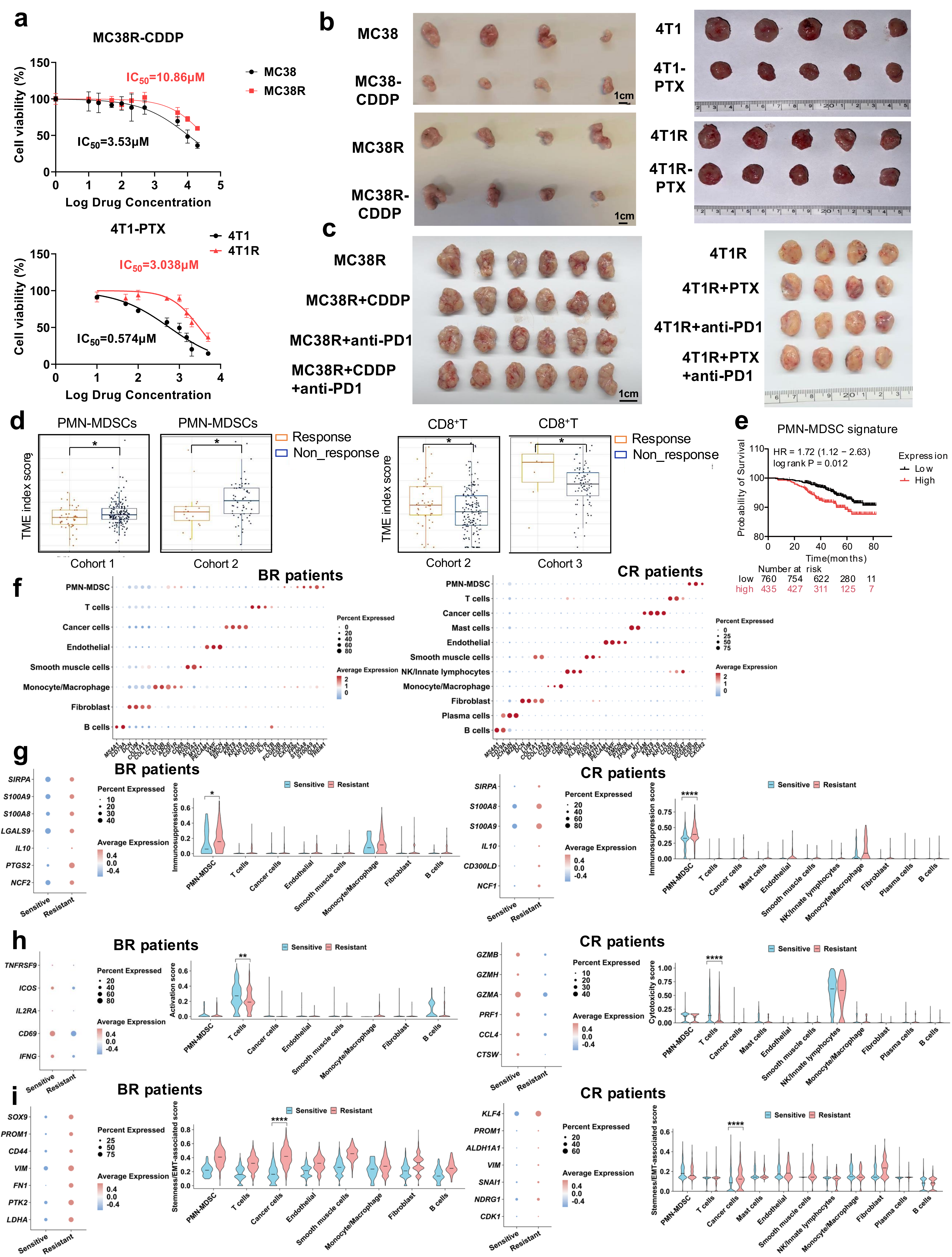
a
MC38R-CDDP
IC50=10.86μM
IC50=3.53μM
MC38
MC38R
Cell viability (%)
Log Drug Concentration
4T1-PTX
4T1
4T1R
IC50=3.038μM
IC50=0.574μM
b
MC38
MC38-CDDP
MC38R
MC38R-CDDP
4T1
4T1-PTX
4T1R
4T1R-PTX
1cm
c
MC38R
MC38R+CDDP
MC38R+anti-PD1
MC38R+CDDP +anti-PD1
4T1R
4T1R+PTX
4T1R+anti-PD1
4T1R+PTX +anti-PD1
d
PMN-MDSCs
CD8+T
Response
Non_response
TME index score
Cohort 1
Cohort 2
Cohort 3
e
PMN-MDSC signature
HR = 1.72 (1.12 − 2.63)
log rank P = 0.012
Expression
Low
High
Probability of Survival
Time(months)
Number at risk
low 760 754 622 280 11
high 435 427 311 125 7
f
BR patients
CR patients
Percent Expressed
Average Expression
g
Sensitive
Resistant
Immunosuppression score
h
Activation score
Cytotoxicity score
i
Stemness/EMT-associated score

**Extended Data Fig.1. Chemoresistant tumors exhibit impaired therapeutic responses and an immunosuppressive microenvironment, related to Fig.1**

**(a)** Drug-response curves of parental MC38 and 4T1 cells and their chemoresistant derivatives. n = 3 biological replicates/group; two-way ANOVA followed by Sidak's multiple-comparisons test.

**(b)** Tumor images from parental or chemoresistant MC38 and 4T1 models with or without chemotherapy. n = 4-5 mice/group. Scale bars, 1 cm.

**(c)** Tumor images from MC38R- and 4T1R-bearing mice receiving chemotherapy and/or anti-PD-1 therapy. MC38R, n = 4-6 mice/group; Scale bars, 1 cm.

**(d)** PMN-MDSC and $CD8^+$T cell gene signature scores in responders and non-responders from independent breast cancer cohorts with neo-adjuvant chemotherapy treatment. n corresponds to the number of patients represented by individual points in each cohort; two-tailed unpaired Student's t test.

**(e)** Kaplan–Meier overall survival according to PMN-MDSC signature level (n = 1,195 patients based on the displayed risk table); log-rank test.

**(f)** scRNA-seq analysis of signature genes for each cell type.

**(g–i)** The differentiated gene expression in each cell type. n represents all cells of the indicated cell populations passing quality control in the corresponding datasets.

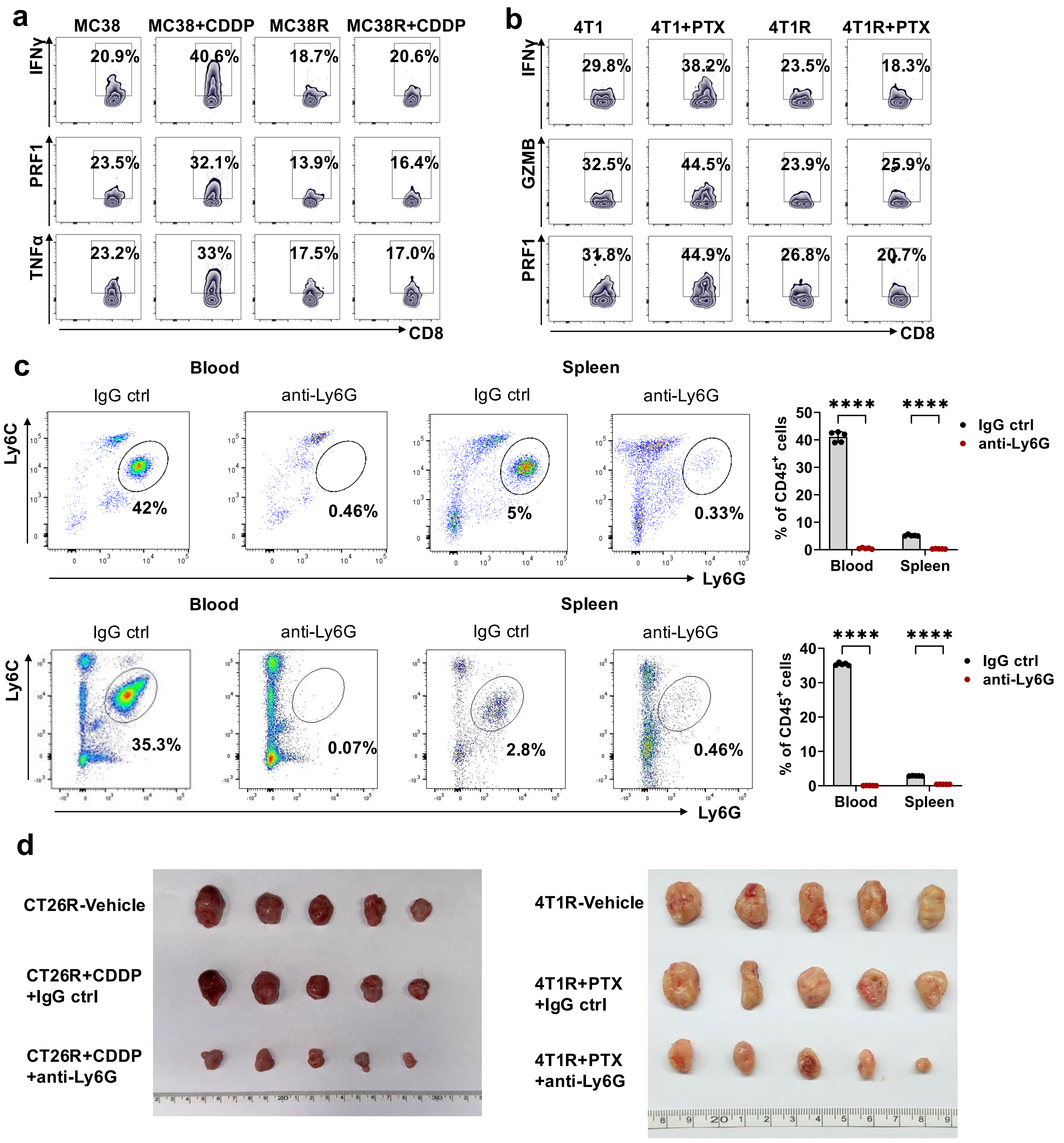
a
MC38
MC38+CDDP
MC38R
MC38R+CDDP
IFNγ
PRF1
TNFα
CD8
20.9%
40.6%
18.7%
20.6%
23.5%
32.1%
13.9%
16.4%
23.2%
33%
17.5%
17.0%
b
4T1
4T1+PTX
4T1R
4T1R+PTX
IFNγ
GZMB
PRF1
CD8
29.8%
38.2%
23.5%
18.3%
32.5%
44.5%
23.9%
25.9%
31.8%
44.9%
26.8%
20.7%
c
Blood
Spleen
IgG ctrl
anti-Ly6G
Ly6C
Ly6G
42%
0.46%
5%
0.33%
35.3%
0.07%
2.8%
0.46%
% of CD45+ cells
****
Blood
Spleen
d
CT26R-Vehicle
CT26R+CDDP +IgG ctrl
CT26R+CDDP +anti-Ly6G
4T1R-Vehicle
4T1R+PTX +IgG ctrl
4T1R+PTX +anti-Ly6G

**Extended Data Fig.2. PMN-MDSCs are essential for mediating chemoresistance, related to Fig. 2**

**(a-b)** Representative flow cytometry plots of cytotoxic effector molecules in $CD8^+$T cells following co-culture with PMN-MDSCs isolated from the indicated tumors. n=5-6 mice/group.

**(c)** Representative flow cytometry plots and quantification of $Ly6G^+$ cells in blood and spleen following control IgG or anti-Ly6G injection. n = 5 mice/group; two-way ANOVA followed by Sidak's multiple-comparisons test.

**(d)** Tumor images of CT26R- and 4T1R-bearing mice following the indicated treatments. n = 5 mice/group.

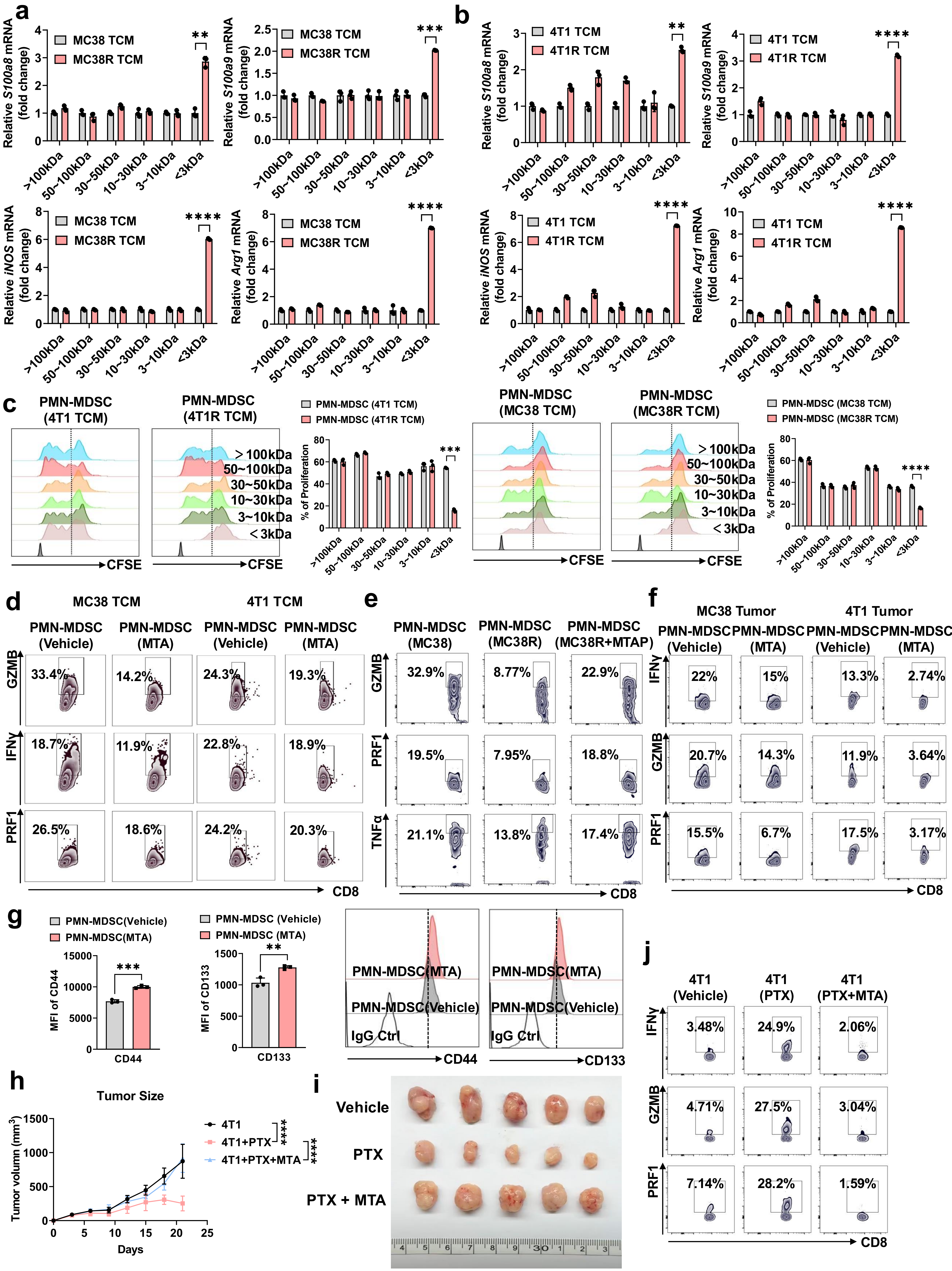
a
MC38 TCM
MC38R TCM
Relative S100a8 mRNA (fold change)
Relative S100a9 mRNA (fold change)
Relative iNOS mRNA (fold change)
Relative Arg1 mRNA (fold change)
>100kDa
50~100kDa
30~50kDa
10~30kDa
3~10kDa
<3kDa
b
4T1 TCM
4T1R TCM
c
PMN-MDSC (4T1 TCM)
PMN-MDSC (4T1R TCM)
PMN-MDSC (MC38 TCM)
PMN-MDSC (MC38R TCM)
% of Proliferation
CFSE
d
MC38 TCM
4T1 TCM
PMN-MDSC (Vehicle)
PMN-MDSC (MTA)
GZMB
IFNγ
PRF1
CD8
e
PMN-MDSC (MC38)
PMN-MDSC (MC38R)
PMN-MDSC (MC38R+MTAP)
TNFα
f
MC38 Tumor
4T1 Tumor
g
PMN-MDSC(Vehicle)
PMN-MDSC(MTA)
MFI of CD44
MFI of CD133
CD44
CD133
IgG Ctrl
h
Tumor Size
Tumor volumn (mm3)
Days
4T1
4T1+PTX
4T1+PTX+MTA
i
Vehicle
PTX
PTX + MTA
j
4T1 (Vehicle)
4T1 (PTX)
4T1 (PTX+MTA)

**Extended Data Fig.3. MTA enhances PMN-MDSC immunosuppressive activity and promotes tumor stemness and chemoresistance, related to Fig. 3**

**(a-b)** Immunosuppressive gene expression in PMN-MDSCs induced by the indicated molecular-weight TCM fractions. n = 3 biological replicates/group; two-way ANOVA followed by Sidak's multiple-comparisons test.

**(c)** $CD8^{+}$T cell proliferation following co-culture with PMN-MDSCs induced by the indicated TCM fractions. n = 3 biological replicates/group; two-way ANOVA followed by Sidak's multiple-comparisons test.

**(d)** Representative flow cytometry plots showing GZMB, IFN-γ, and PRF1 expression in $CD8^{+}$T cells following co-culture with vehicle- or MTA-treated PMN-MDSCs induced by parental MC38 or 4T1 TCM, corresponding to the quantitative analysis shown in Fig. 3e. Representative of n = 3-6 biological replicates/group.

**(e)** Representative flow cytometry plots showing GZMB, PRF1, and TNF-α expression in $CD8^{+}$T cells following co-culture with PMN-MDSCs induced by MC38 TCM, MC38R TCM, or MC38R TCM treated with MTAP, corresponding to the quantitative analysis shown in Fig. 3f. Representative of n = 3 biological replicates/group.

**(f)** Representative flow cytometry plots showing IFN-γ, GZMB, and PRF1 expression in $CD8^{+}$T cells following co-culture with vehicle- or MTA-treated PMN-MDSCs isolated from MC38 or 4T1 tumors, corresponding to the quantitative analysis shown in Fig. 3h. Representative of n = 5 biological replicates/group.

**(g)** CD44 and CD133 expression in 4T1 tumor cells following co-culture with vehicle- or MTA-treated PMN-MDSCs. n = 3 biological replicates/group; two-tailed unpaired Student's t test for each marker.

**(h)** 4T1 tumor growth following vehicle, PTX, or PTX plus MTA. n = 5 mice/group; repeated-measures two-way ANOVA followed by Sidak's multiple-comparisons test.

**(i)** Tumor images of 4T1 as described in (h). n = 5 mice/group.

**(j)** Representative flow cytometry plots showing IFN-γ, GZMB, and PRF1 expression in $CD8^{+}$T cells following co-culture with PMN-MDSCs isolated from tumors as described in (h). n = 5 mice/group.

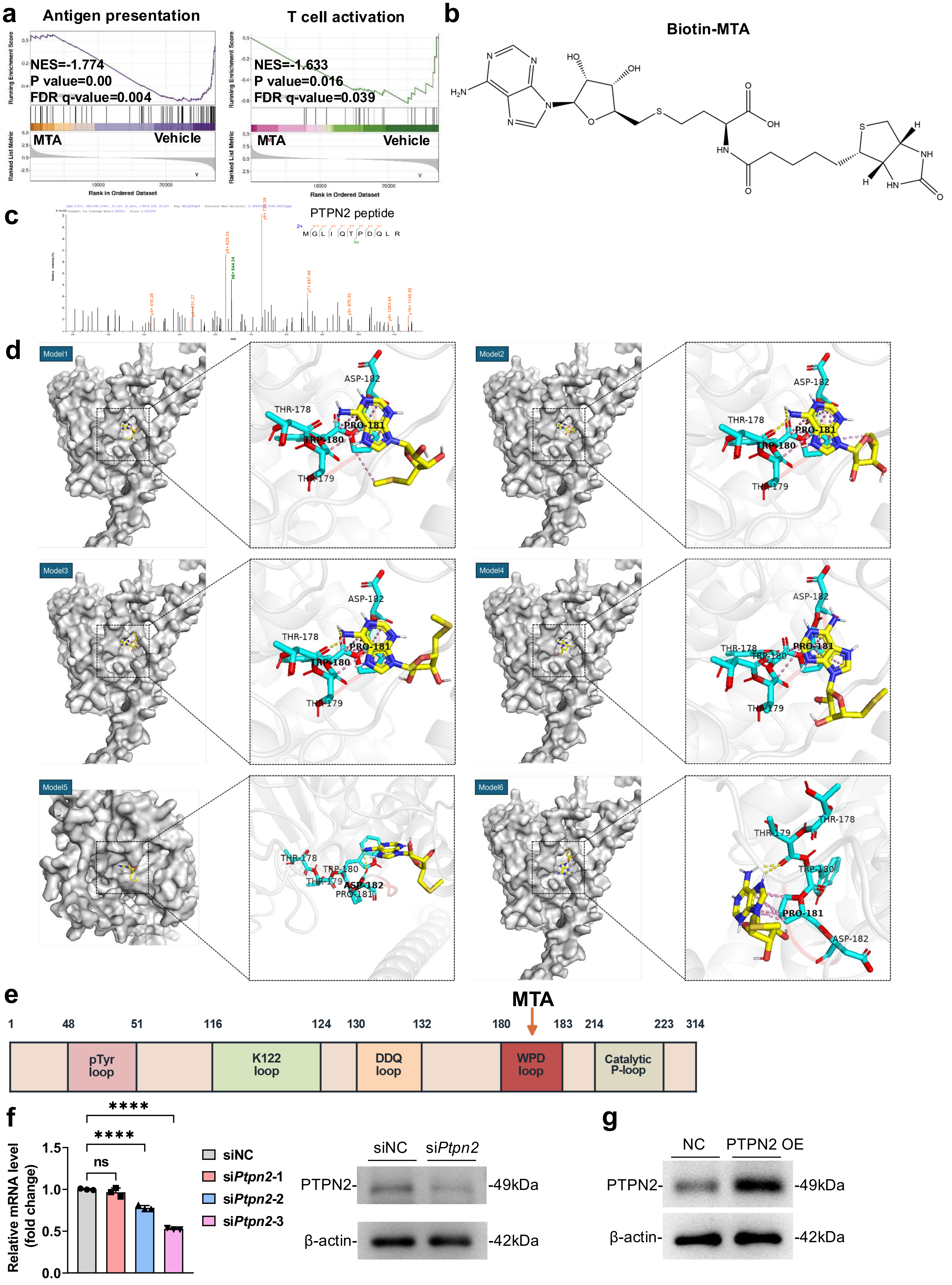

a
Antigen presentation
NES=-1.774
P value=0.00
FDR q-value=0.004
MTA
Vehicle
T cell activation
NES=-1.633
P value=0.016
FDR q-value=0.039
MTA
Vehicle
Running Enrichment Score
Ranked List Metric
Rank in Ordered Dataset
b
Biotin-MTA
c
PTPN2 peptide
M G L I Q T P D Q L R
d
Model1
Model2
Model3
Model4
Model5
Model6
ASP-182
THR-178
PRO-181
TRP-180
THR-179
e
MTA
1 48 51 116 124 130 132 180 183 214 223 314
pTyr loop
K122 loop
DDQ loop
WPD loop
Catalytic P-loop
f
Relative mRNA level (fold change)
ns
****
****
siNC
siPtpn2-1
siPtpn2-2
siPtpn2-3
siNC
siPtpn2
PTPN2-
-49kDa
β-actin-
-42kDa
g
NC
PTPN2 OE
PTPN2-
-49kDa
β-actin-
-42kDa

**Extended Data Fig. 4. MTA associates with the WPD-loop region of PTPN2 and regulates PTPN2-dependent JAK–STAT3 signaling, related to Fig. 4**

**(a)** GSEA analysis of immune-related genes in vehicle- and MTA-treated PMN-MDSCs, n = 3 biological replicates/group.

**(b)** Chemical structure of biotin–MTA.

**(c)** LC–MS/MS spectrum of a PTPN2-derived peptide identified in the biotin–MTA affinity-purification sample.

**(d)** Six predicted docking poses of MTA with PTPN2 , n = 6 computational models.

**(e)** Schematic illustration of PTPN2 functional domains.

**(f)** RT-qPCR and Western blot analysis of PTPN2 expression in PMN-MDSCs following *Ptpn2* siRNA transfection. n = 3 biological samples; one-way ANOVA followed by Sidak's multiple-comparisons test. Immunoblot is representative of three independent experiments.

**(g)** Western blot analysis of PTPN2 expression in PMN-MDSCs after electroporation with PTPN2 overexpression plasmid (pLV-CMV-EGFP-m*Ptpn2*) ; representative of three independent experiments.

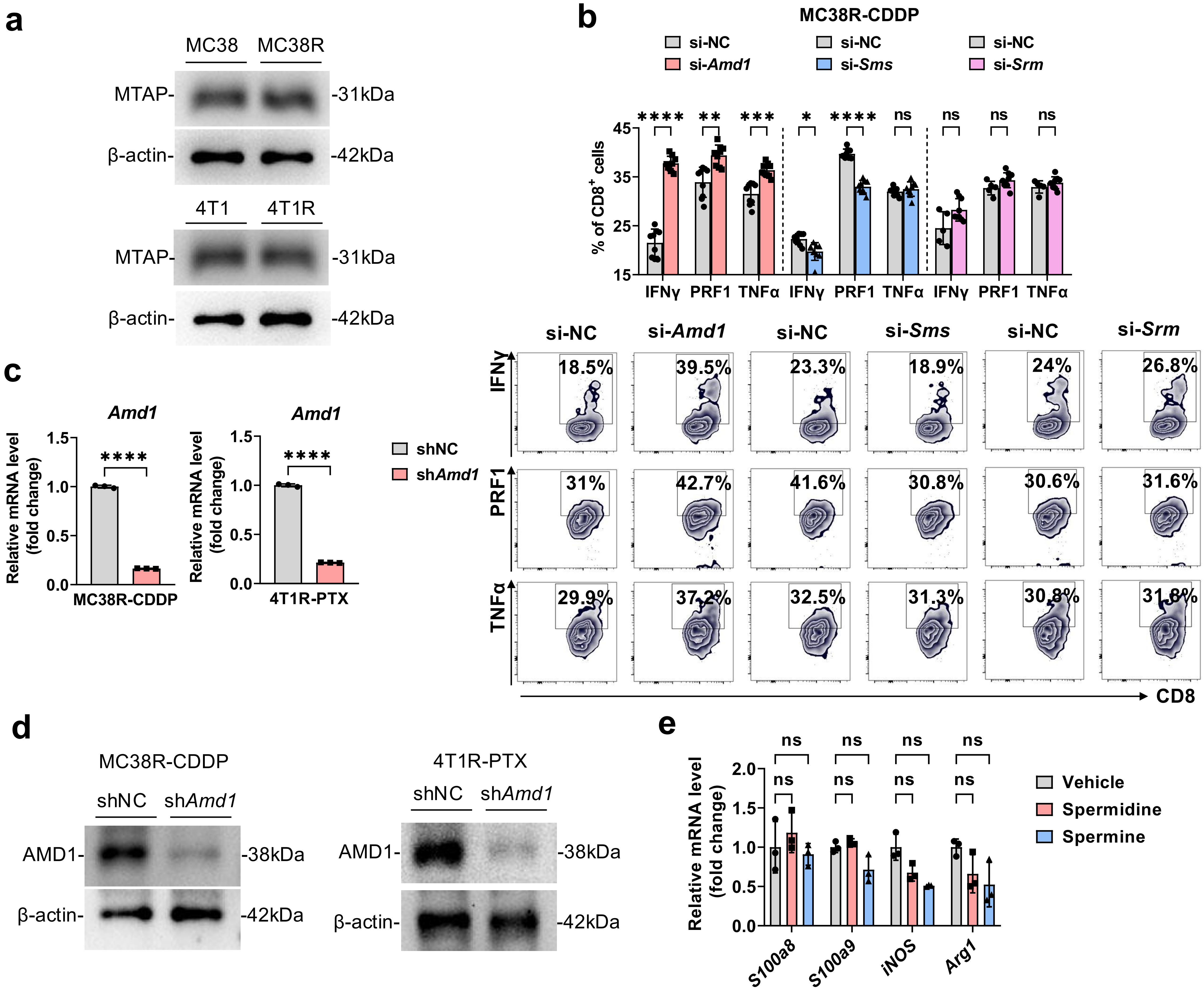
a
MC38
MC38R
MTAP-
-31kDa
β-actin-
-42kDa
4T1
4T1R
MTAP-
-31kDa
β-actin-
-42kDa
b
MC38R-CDDP
si-NC
si-Amd1
si-NC
si-Sms
si-NC
si-Srm
% of CD8+ cells
IFNγ
PRF1
TNFα
si-NC
si-Amd1
si-NC
si-Sms
si-NC
si-Srm
CD8
c
Amd1
Relative mRNA level (fold change)
MC38R-CDDP
4T1R-PTX
shNC
shAmd1
d
MC38R-CDDP
shNC
shAmd1
AMD1-
-38kDa
β-actin-
-42kDa
4T1R-PTX
shNC
shAmd1
AMD1-
-38kDa
β-actin-
-42kDa
e
Relative mRNA level (fold change)
S100a8
S100a9
iNOS
Arg1
Vehicle
Spermidine
Spermine

**Extended Data Fig.5. AMD1 is required for MTA-mediate PMN-MDSC immunosuppressive activity, related to Fig. 5**

**(a)** MTAP immunoblots of parental and chemoresistance tumor cells; representative of three independent experiments.

**(b)** Quantification and representative flow cytometry plots of IFN-γ, PRF1, and TNF-α expression in $CD8^{+}$T cells following co-culture with PMN-MDSCs induced by TCM from MC38R cells transfected with control siRNA or siRNAs targeting *Amd1*, *Sms*, or *Srm*. n = 5–9 biological replicates/group. Data are shown as mean ± SEM; two-way ANOVA followed by Sidak's multiple-comparisons test.

**(c)** RT–qPCR validation of *Amd1* knockdown in chemoresistant tumor cells, n = 3 biological replicates; two-tailed unpaired Student's t test.

**(d)** Immunoblot validation of AMD1 knockdown in chemoresistant tumor cells; representative of three independent experiments.

**(e)** Immunosuppressive gene expression following vehicle, spermidine, or spermine treatment. n = 3 biological replicates; two-way ANOVA followed by Sidak's multiple-comparisons test.

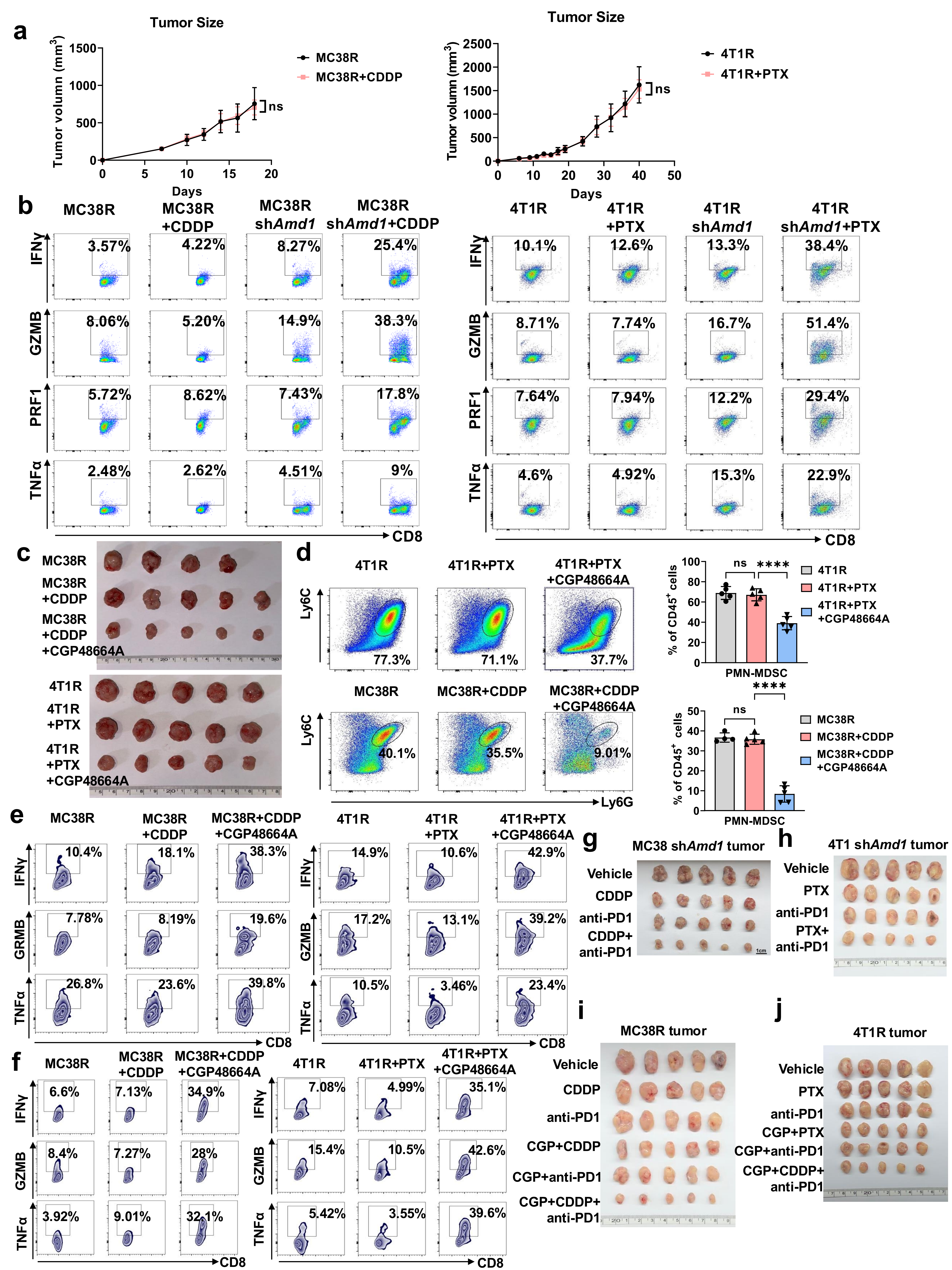

a
Tumor Size
Tumor volumn (mm3)
Days
MC38R
MC38R+CDDP
ns
4T1R
4T1R+PTX
b
MC38R
MC38R +CDDP
MC38R shAmd1
MC38R shAmd1+CDDP
4T1R
4T1R +PTX
4T1R shAmd1
4T1R shAmd1+PTX
IFNγ
GZMB
PRF1
TNFα
CD8
c
MC38R
MC38R +CDDP
MC38R +CDDP +CGP48664A
4T1R
4T1R +PTX
4T1R +PTX +CGP48664A
d
4T1R
4T1R+PTX
4T1R+PTX +CGP48664A
Ly6C
Ly6G
MC38R
MC38R+CDDP
MC38R+CDDP +CGP48664A
% of CD45+ cells
PMN-MDSC
ns
****
e
MC38R
MC38R +CDDP
MC38R+CDDP +CGP48664A
4T1R
4T1R +PTX
4T1R+PTX +CGP48664A
IFNγ
GRMB
GZMB
TNFα
CD8
f
g
MC38 shAmd1 tumor
Vehicle
CDDP
anti-PD1
CDDP+ anti-PD1
h
4T1 shAmd1 tumor
Vehicle
PTX
anti-PD1
PTX+ anti-PD1
i
MC38R tumor
Vehicle
CDDP
anti-PD1
CGP+CDDP
CGP+anti-PD1
CGP+CDDP+ anti-PD1
j
4T1R tumor
Vehicle
PTX
anti-PD1
CGP+PTX
CGP+anti-PD1
CGP+CDDP+ anti-PD1

**Extended Data Fig. 6. AMD1 inhibition restores therapeutic responsiveness and alleviates PMN-MDSC-mediated immunosuppression, related to Fig. 6**

**(a)** Tumor growth of MC38R and 4T1R tumors with or without chemotherapy. n = 5 mice/group; repeated-measures two-way ANOVA followed by Sidak's multiple-comparisons test.

**(b)** Representative flow cytometry plots showing IFN-γ, GZMB, PRF1, and TNF-α expression in $CD8^+$T cells following co-culture with PMN-MDSCs isolated from control or *Amd1*-knockdown MC38R and 4T1R tumors with or without the corresponding chemotherapy, corresponding to the quantitative analysis shown in Fig. 6b. Representative of n = 5 mice/group.

**(c)** Tumor images of MC38R and 4T1R tumors following chemotherapy treatment, or combined with CGP48664A. n = 5 mice/group.

**(d)** The percentages of intratumoral PMN-MDSC in MC38R and 4T1R tumors following chemotherapy treatment, or combined with CGP48664A. n = 5 mice/group; one-way ANOVA followed by Sidak's multiple-comparisons test.

**(e)** Representative flow cytometry plots showing IFN-γ, GZMB, and TNF-α expression in tumor-infiltrating $CD8^+$T cells from MC38R and 4T1R tumors following chemotherapy treatment, or combined with CGP48664A, corresponding to the quantitative analysis shown in Fig. 6e. Representative of n = 5 mice/group.

**(f)** Representative flow cytometry plots showing IFN-γ, GZMB, and TNF-α expression in $CD8^+$T cells following ex vivo co-culture with PMN-MDSCs isolated from the indicated treatment groups, corresponding to the quantitative analysis shown in Fig. 6f. Representative of n = 5 biological replicates/group.

**(g–j)** Tumor images from the indicated treatment groups, n = 5 mice/group.

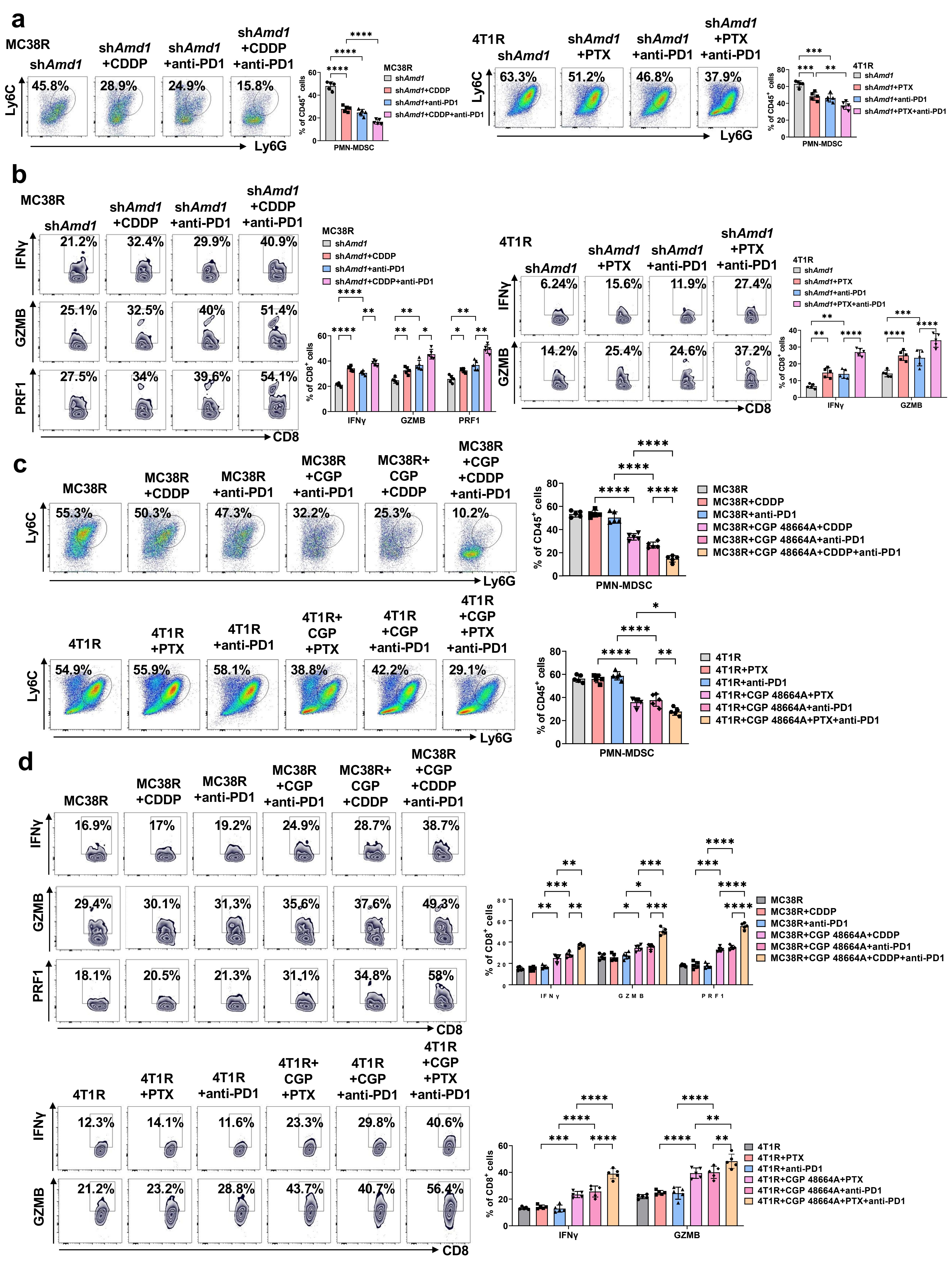
a
MC38R
shAmd1
shAmd1 +CDDP
shAmd1 +anti-PD1
shAmd1 +CDDP +anti-PD1
PMN-MDSC
4T1R
shAmd1 +PTX
shAmd1 +anti-PD1
shAmd1 +PTX +anti-PD1
Ly6C
Ly6G
% of CD45+ cells
b
IFNγ
GZMB
PRF1
CD8
% of CD8+ cells
c
MC38R+CDDP
MC38R+anti-PD1
MC38R+CGP 48664A+CDDP
MC38R+CGP 48664A+anti-PD1
MC38R+CGP 48664A+CDDP+anti-PD1
4T1R+PTX
4T1R+anti-PD1
4T1R+CGP 48664A+PTX
4T1R+CGP 48664A+anti-PD1
4T1R+CGP 48664A+PTX+anti-PD1
d

**Extended Data Fig. 7. AMD1 inhibition remodels the tumor immune microenvironment during chemo-immunotherapy, related to Fig. 6**

**(a)** The percentages of intratumoral PMN-MDSCs in *Amd1*-knockdown tumors following chemotherapy and/or anti-PD-1 treatment. n = 5 mice/group; one-way ANOVA followed by Sidak's multiple-comparisons test.

**(b)** The effector molecule expression of tumor-infiltrating $CD8^{+}$T cells in the indicated tumors as described in (a). n = 5 mice/group; two-way ANOVA followed by Sidak's multiple-comparisons test.

**(c)** The percentages of intratumoral PMN-MDSCs following pharmacological AMD1 inhibition with chemotherapy and/or anti-PD-1 treatment. n = 5 mice/group; one-way ANOVA followed by Sidak's multiple-comparisons test.

**(d)** The effector molecule expression of tumor-infiltrating $CD8^{+}$T cell in the indicated tumors as described in (c). n = 5 mice/group; two-way ANOVA followed by Sidak's multiple-comparisons test.

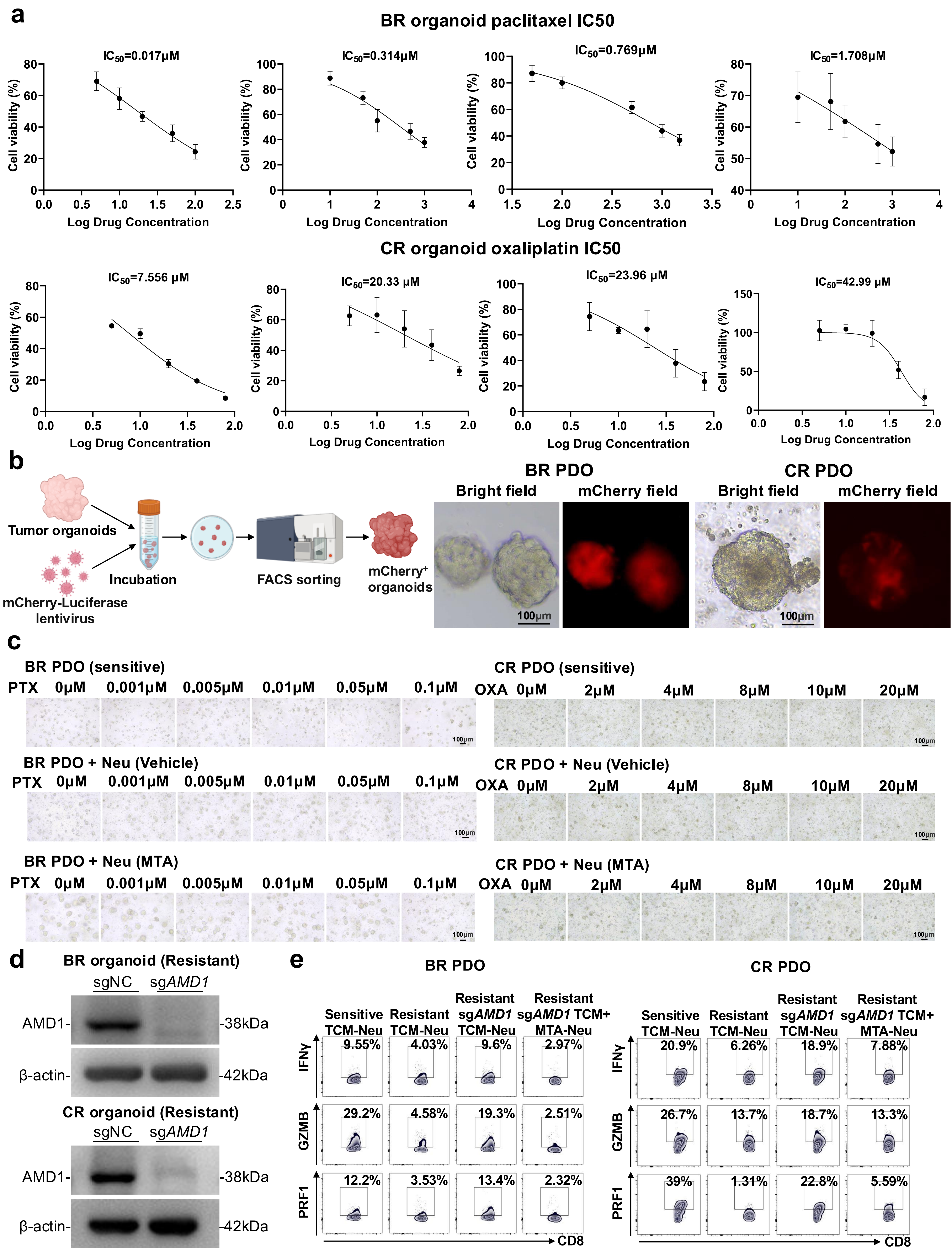

a
BR organoid paclitaxel IC50
IC50=0.017μM
IC50=0.314μM
IC50=0.769μM
IC50=1.708μM
Cell viability (%)
Log Drug Concentration
CR organoid oxaliplatin IC50
IC50=7.556 μM
IC50=20.33 μM
IC50=23.96 μM
IC50=42.99 μM
b
Tumor organoids
mCherry-Luciferase lentivirus
Incubation
FACS sorting
mCherry+ organoids
BR PDO
CR PDO
Bright field
mCherry field
100μm
c
BR PDO (sensitive)
PTX 0μM 0.001μM 0.005μM 0.01μM 0.05μM 0.1μM
CR PDO (sensitive)
OXA 0μM 2μM 4μM 8μM 10μM 20μM
BR PDO + Neu (Vehicle)
CR PDO + Neu (Vehicle)
BR PDO + Neu (MTA)
CR PDO + Neu (MTA)
d
BR organoid (Resistant)
sgNC sgAMD1
AMD1- -38kDa
β-actin- -42kDa
CR organoid (Resistant)
e
BR PDO
CR PDO
Sensitive TCM-Neu
Resistant TCM-Neu
Resistant sgAMD1 TCM-Neu
Resistant sgAMD1 TCM+ MTA-Neu
IFNγ
GZMB
PRF1
CD8
9.55% 4.03% 9.6% 2.97%
29.2% 4.58% 19.3% 2.51%
12.2% 3.53% 13.4% 2.32%
20.9% 6.26% 18.9% 7.88%
26.7% 13.7% 18.7% 13.3%
39% 1.31% 22.8% 5.59%

**Extended Data Fig. 8. Establishment and functional characterization of patient-derived tumor organoid models, related to Fig. 7**

**(a)** Chemotherapeutic drug-response curves and IC50 values of individual breast and colorectal cancer PDOs. n = 3 biological replicates.

**(b)** mCherry–luciferase PDO establishment. Images are representative of three biological replicates/group. Scale bars, 100 μm.

**(c)** Representative bright-field images of PDOs co-cultured with vehicle- or MTA-treated neutrophils following chemotherapeutic drug treatment. Scale bars, 100 μm.

**(d)** AMD1 depletion in chemoresistant PDOs by CRISPR–Cas9; immunoblots are representative of three independent experiments.

**(e)** Representative flow cytometry plots showing IFN-γ, GZMB, and PRF1 expression in human $CD8^{+}$T cells following co-culture with healthy donor-derived neutrophils preconditioned with media from chemosensitive, chemoresistant, AMD1-deficient chemoresistant, or AMD1-deficient chemoresistant breast cancer and colorectal cancer PDOs with MTA addition, corresponding to the quantitative analysis shown in Fig. 7g. Representative of n = 5 biological replicates.

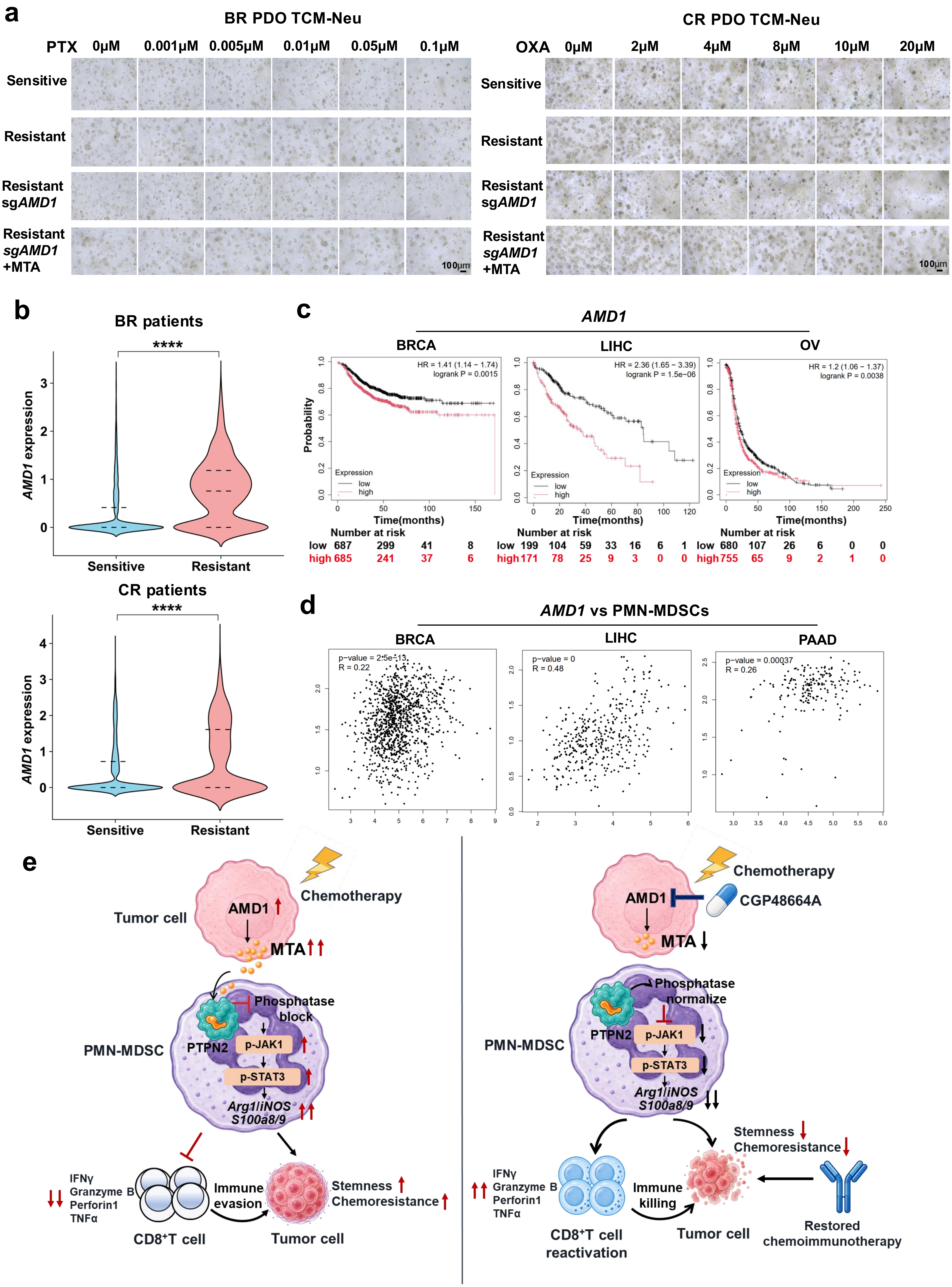
a
BR PDO TCM-Neu
PTX 0μM 0.001μM 0.005μM 0.01μM 0.05μM 0.1μM
Sensitive
Resistant
Resistant sgAMD1
Resistant sgAMD1 +MTA
100μm
CR PDO TCM-Neu
OXA 0μM 2μM 4μM 8μM 10μM 20μM
b
BR patients
CR patients
AMD1 expression
Sensitive
Resistant
****
c
AMD1
BRCA
LIHC
OV
HR = 1.41 (1.14 − 1.74) logrank P = 0.0015
HR = 2.36 (1.65 − 3.39) logrank P = 1.5e−06
HR = 1.2 (1.06 − 1.37) logrank P = 0.0038
Probability
Expression
low
high
Time(months)
Number at risk
low 687 299 41 8
high 685 241 37 6
low 199 104 59 33 16 6 1
high 171 78 25 9 3 0 0
low 680 107 26 6 0 0
high 755 65 9 2 1 0
d
AMD1 vs PMN-MDSCs
BRCA
LIHC
PAAD
p-value = 2.5e-13
R = 0.22
p-value = 0
R = 0.48
p-value = 0.00037
R = 0.26
e
Tumor cell
AMD1
Chemotherapy
MTA
PMN-MDSC
PTPN2
Phosphatase block
p-JAK1
p-STAT3
Arg1/iNOS
S100a8/9
IFNγ
Granzyme B
Perforin1
TNFα
CD8⁺T cell
Immune evasion
Stemness
Chemoresistance
CGP48664A
Phosphatase normalize
CD8⁺T cell reactivation
Immune killing
Restored chemoimmunotherapy

**Extended Data Fig. 9. Clinical relevance and working model of the AMD1–MTA–PMN-MDSC axis, related to Fig. 7**

**(a)** Representative PDO images corresponding to Fig.7h. Scale bars, 100 μm.

**(b)** AMD1 expression in the tumor cells of chemo-immunotherapy sensitive and resistant patients based on the scRNA-seq data (GSE302453 and GSE236581).

**(c)** Kaplan–Meier survival analyses according to AMD1 expression: BRCA, n = 1,372; LIHC, n = 370; OV, n = 1,435, based on the displayed numbers at risk; log-rank test.

**(d)** Correlations between AMD1 expression and PMN-MDSC signature scores in BRCA, LIHC, and PAAD.

**(e)** Proposed working model.
