## Supplementary material for "AMD1-driven MTA production sustains chemoresistance by rewiring PMN-MDSC immunosuppression": Table S1

### Table S1. Nucleotide sequences

| **qPCR Primers** | **Sequence (5’-3’)** |
|---|---|
| Mouse *S100a8* qPCR forward primer | AAATCACCATGCCCTCTACAAG |
| Mouse *S100a8* qPCR reverse primer | CCCACTTTTATCACCATCGCAA |
| Mouse *S100a9* qPCR forward primer | ATACTCTAGGAAGGAAGGACACC |
| Mouse *S100a9* qPCR reverse primer | TCCATGATGTCATTTATGAGGGC |
| Mouse *Arg1* qPCR forward primer | GGCAACCTGTGTCCTTTCTCCTG |
| Mouse *Arg1* qPCR reverse primer | GGTCTACGTCTCGCAAGCCAATG |
| Mouse *Nos2* qPCR forward primer | ACTCAGCCAAGCCCTCACCTAC |
| Mouse *Nos2* qPCR reverse primer | TCCAATCTCTGCCTATCCGTCTCG |
| Mouse *Amd1* qPCR forward primer | TCAGTAAGTGCTGCCGTTGCCA |
| Mouse *Amd1* qPCR reverse primer | AGACCAGAGACCAGCTGACGGT |
| Human *AMD1* qPCR forward primer | CTGGTTCCCCTGTTGAAGCT |
| Human *AMD1* qPCR reverse primer | GGTACCCTTGGTGAGAAGGC |
| Human *AMD1* editing-locus forward primer | CAGCTATAGGCCGTGGAGGT |
| Human *AMD1* editing-locus reverse primer | GCCGCTGCGGATTGTTCCAG |
| **CRISPR sgRNA** | |
| Human *AMD1* sgRNA cloning oligo, forward | CACCGCGAGCGCTAGCTAATGTCGC |
| Human *AMD1* sgRNA cloning oligo, reverse | AAACGCGACATTAGCTAGCGCTCGC |
| **Mouse PTPN2 siRNAs** | |
| Mus-*Ptpn2*-siR1 sense | CCAGCUUAGUUGACAUAGAdTdT |
| Mus-*Ptpn2*-siR1 antisense | UCUAUGUCAACUAAGCUGGdTdT |
| Mus-*Ptpn2*-siR2 sense | GAGAAUAGGUUCAGAAGAUdTdT |
| Mus-*Ptpn2*-siR2 antisense | AUCUUCUGAACCUAUUCUCdTdT |
| Mus-*Ptpn2*-siR3 sense | GUAGAAAGAUGUAAAGGAAdTdT |
| Mus-*Ptpn2*-siR3 antisense | UUCCUUUACAUCUUUCUACdTdT |
| **Mouse SRM siRNAs** | |
| *Srm*-681-s | CAUGAAGACAGCACUCAAAdTdT |
| *Srm*-681-a | UUUGAGUGCUGUCUUCAUGdTdT |
| *Srm*-542-s | GCUCAAAGCUGACUCUCCAdTdT |
| *Srm*-542-a | UGGAGAGUCAGCUUUGAGCdTdT |
| *Srm*-386-s | CGCGGAAGGUGCUGAUCAUdTdT |
| *Srm*-386-a | AUGAUCAGCACCUUCCGCGdTdT |
| **Mouse SMS siRNAs** | |
| *Sms*-419-s | GGUUGAGUAUGACAUAGAUdTdT |
| *Sms*-419-a | AUCUAUGUCAUACUCAACCdTdT |
| Sms-736-s | GUGGAGAUGUCUUAGACAAdTdT |
| Sms-736-a | UUGUCUAAGACAUCUCCACdTdT |
| *Sms*-1108-s | GGAAGAAAGCUAAGCCUUAdTdT |
| Sms-1108-a | UAAGGCUUAGCUUUCUUCCdTdT |
| **Mouse AMD1 siRNAs** | |
| *Amd1*-895-s | GGACCAGAUCUACAUGAAAdTdT |
| *Amd1*-895-a | UUUCAUGUAGAACUGGUCCdTdT |
| *Amd1*-1134-s | GGAAAGUUGUGGAAGUCUUdTdT |
| *Amd1*-1134-a | AAGACUUCCACAACUUUCCdTdT |
| *Amd1*-477-s | CAAUCAUAAGUGUGACAAAdTdT |
| *Amd1*-477-a | UUUGUCACACUUAUGAUUGdTdT |
| **Mouse *Amd1* shRNA** | |
| *Amd1* shRNA sequence | TACAGTGGGTTTGACTCAATT |
